# Experimental evolution of adaptive divergence under varying degrees of gene flow

**DOI:** 10.1101/2020.11.02.364695

**Authors:** Sergio Tusso, Bart P.S. Nieuwenhuis, Bernadette Weissensteiner, Simone Immler, Jochen B.W. Wolf

**Author notes:** Corresponding authors: S. Tusso:, J. B.W. Wolf. These authors contributed equally: S. Tusso, B. Nieuwenhuis. These authors jointly supervised this work: S. Immler, J. Wolf.

## Abstract

Adaptive divergence is the key evolutionary process generating biodiversity by means of natural selection. Yet, the conditions under which it can arise in the presence of gene flow remain contentious. To address this question, we subjected 132 sexually reproducing fission yeast populations sourced from two independent genetic backgrounds to disruptive ecological selection and manipulated the level of migration between environments. Contrary to theoretical expectations, adaptive divergence was most pronounced when migration was either absent (‘allopatry’) or maximal (‘sympatry’), but was much reduced at intermediate rates (‘parapatry’, ‘local mating’). This effect was apparent across central life history components (survival, asexual growth, and mating), but differed in magnitude between ancestral genetic backgrounds. The evolution of some fitness components was constrained by pervasive negative correlations (trade-off between asexual growth and mating), while others changed direction under the influence of migration (e.g. survival and mating). In allopatry, adaptive divergence was mainly conferred by standing genetic variation and resulted in ecological specialization. In sympatry, divergence was mainly mediated by novel mutations enriched in a subset of genes and was characterized by the repeated emergence of two strategies: an ecological generalist and an asexual growth specialist. Multiple loci showed consistent evidence for antagonistic pleiotropy across migration treatments and provide a conceptual link between adaptation and divergence. This evolve-and-resequence experiment demonstrates that rapid ecological differentiation can arise even under high rates of gene flow. It further highlights that adaptive trajectories are governed by complex interactions of gene flow, ancestral variation and genetic correlations.

## Main

Adaptive divergence describes the emergence of new forms from a shared common ancestor by adaptation to different environmental conditions. As such, it is key to the formation of new species by means of natural selection ^1,2^. In geographic isolation, divergent selection readily promotes ecological specialisation which over time can result in reproductive barriers between populations ^3–6^. In the presence of gene flow, however, the conditions enabling adaptive divergence are difficult to predict ^7–9^. Homogenizing gene flow may impede adaptive divergence and promote generalist phenotypes exploiting a broader ecological spectrum ^10,11^. Alternatively, gene flow may promote divergence by supplying adaptive genetic variation or by modifying genetic correlations which can alter evolutionary constraints and open new evolutionary trajectories ^12,13^. In general, the relationship between gene flow and adaptive divergence is expected to decline monotonically (i.e. less divergence with higher gene flow) ^14^. Depending on the degree of gene flow, the strength of selection, and the genetic architecture of adaptive traits evolutionary outcomes are, however, hard to predict: divergence may be precluded, stalled at an intermediate level or progress towards the origin of new species ^15–17,14^. Adaptive divergence thus constitutes a necessary, but not sufficient component for ecological speciation. Understanding the conditions under which it arises is of central importance to our understanding of how populations can exploit divergent ecological niches and differentiate into distinct ecotypes that may – or may not – seed novel species ^18,19^.

Genome-wide characterization of genetic variation has spurred progress in the study of ecological divergence with gene flow in the wild ^20–22^. However, the idiosyncratic nature and complex evolutionary histories of natural populations impair inference of causal relationships and make it difficult to pinpoint the mechanisms promoting or impeding divergence ^8^. Controlled experiments elucidating the genetic basis of adaptive divergence and evaluating the role of gene flow are thus needed ^23,20,24^. Replicated experimental manipulation of migration between controlled ecological contrasts in evolving populations are a promising, although hitherto largely unexplored way forward ^23^.

Here, we present the results from a long-term experimental evolution study addressing this question in the haploid fission yeast *Schizosaccharomyces pombe*, in which we tested the effect of gene flow and standing variation on genetic and phenotypic adaptation to disruptive selection. A total of 132 populations were maintained for 53 complete reproductive cycles each encompassing ∼13 asexual cell divisions. Each cycle comprised asexual growth, followed by ecologically disruptive selection and subsequent sexual reproduction (**Figure 1a**). Sets of 22 populations were distributed among four treatment groups varying in the amount of migration after disruptive selection (ranging from complete isolation to full mixing; hereafter referred to as ‘allopatry’, ‘parapatry’, ‘local mating’, and ‘sympatry’, **Figure 1b & Supplementary Figure 1**). As an ecological parameter we used disruptive viability selection on settling speed by collecting cells from the bottom (bottom selection - B) or the top (top selection - T) in a liquid column after a predefined period of time. Population sizes were in the order of ∼3·10^7^ individuals precluding a dominant role of genetic drift. All experimental populations were derived from two ancestral populations (referred as ‘α’ and ‘β’) that had experienced the same selection regime in the past but differed in standing genetic variation (see **Methods**). For ease of presentation, we will focus on the results for the α genetic background which in general showed a stronger response to selection. We refer to the β background where it deviates from the α. After 53 cycles of sexual reproduction, we measured fitness relative to the respective ancestral population for four fitness components reflecting major life history traits: asexual growth rate (*g*), reproductive success (*r*) and survival during top or bottom selection (top: *W*_*T*_; or bottom: *W*_*B*_).

**Figure 1.**
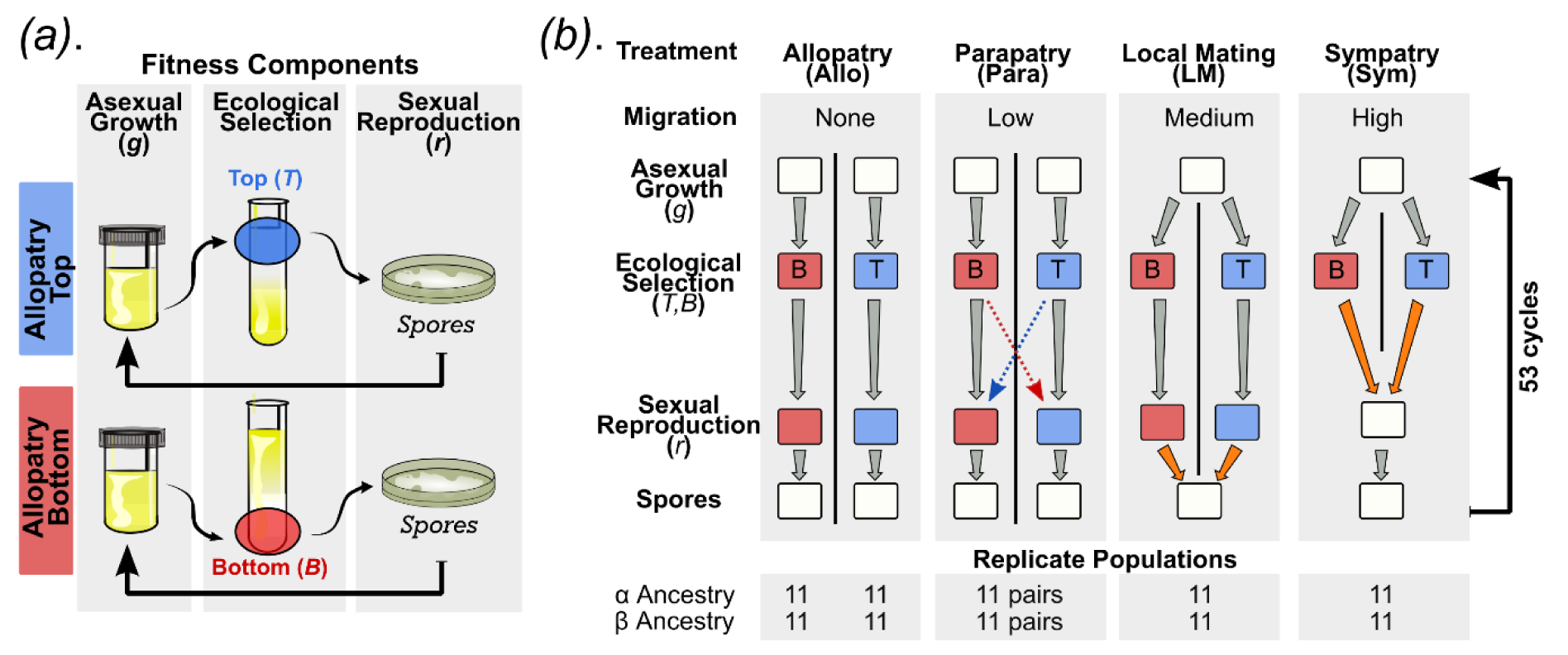
Schematic illustration of the experiment. **a**. Schematic of a 6-day experimental cycle including asexual population growth, ecological selection (top and bottom selection regime) and sexual reproduction (allopatry treatment shown as example). **b**. Representation of the four treatments differing in the amount of migration between fractions after ecological selection: i) allopatry: no gene flow, ii) parapatry: symmetric migration of 5% of cells (red and blue dashed lines), and full mixing (orange arrows) either iii) after sexual reproduction (‘local mating’) or iv) before sexual reproduction (sympatry). The number of populations per treatment for the α or β ancestral background is given at the bottom.

This experiment tests the role of strong divergent ecological selection at four levels of migration. Despite the apparent simplicity of the setup, the experimental life cycle involves a variety of fitness components each of which can be subject to selection, and hence can evolve (e.g. alternation between liquid and solid media, asexual growth, sexual reproduction, survival during ecological selection). Interdependence between fitness components is expected to elicit a correlated response which can promote or constrain adaptive divergence ^25,12^. For instance, increased performance during (asexual) growth can result in reduced output during sexual reproduction ^26–29^. In addition, ecological adaptation may not result even if a given trait value increases survival, but has negative consequences on a correlated life history trait. Importantly, these correlations need not be static as evolution of one life history trait may alter the strength and direction of selection on another (e.g. differences in asexual growth rate bears on population density and thus nutrient availability). Genetically encoded variance-covariance relationships between life history traits (as represented by the G-matrix ^30,31^), and the ability of these relationships to evolve in themselves constitutes an essential component determining the evolutionary trajectory of each population ^13^. We anticipate that the influence and stability of these correlations may be contingent on the level of gene flow and play a central role in constraining or facilitating divergence ^12^.

We report the results of this experiment as follows. We first consider the influence of migration and standing genetic variation on adaptive divergence of each fitness component in isolation. In brief, we found that the degree of adaptive divergence overall depended on standing genetic variation, and was strongest at the extreme ends of the gene flow gradient (allopatry and sympatry). We then expand on these results considering the intrinsic correlations between fitness components (G-matrix) which evolved relative to the ancestral populations in response to gene flow and played a central role in facilitating adaptive divergence. Finally, we assess the genetic architecture of divergence by means of whole genome sequencing data for all ancestral and evolved populations (Pool-Seq). Standing genetic variation with evidence for antagonistic pleiotropy was the main driver for ecological specialization in the absence of gene flow. In contrast, divergence in sympatry was characterized by population-specific independent mutations.

## Results and discussion

### Migration affects adaptive divergence

After 53 cycles of disruptive ecological selection populations showed evidence for evolution across different fitness components (relative fitness difference from ancestral value of 1, see **Figure 2a**). In line with theoretical predictions that isolated populations will readily respond to directional selection^11^, allopatric populations of the α background showed evidence for adaptive divergence in three of the four fitness components (*g, r* and *W*_*T*_) resulting in specialized top and bottom ecotypes (**Figure 2a, Supplementary Table 1 & 2**). Populations that had exclusively been selected for the top environment (shown in blue across figures) grew faster and survived significantly better during further ecological top selection than populations that had only experienced the bottom environment (shown in red). However, their sexual reproductive success was reduced in comparison to bottom populations. These differences were consistent between populations demonstrating that the disruptive ecological selection regime predictably induced the evolution of specialized strategies for the top and bottom environment (for statistical model see **Supplementary Tables 1 & 2)**. Similar divergence across fitness components was observed for the β background, although they showed an overall weaker response and evolved differences in survival after bottom selection rather than for asexual growth. This difference may be related to a diminishing-return relationship for the already higher growth rate of the β ancestor reducing the potential for adaptation in this trait ^32^ (**Supplementary Fig. 2**).

**Figure 2.**
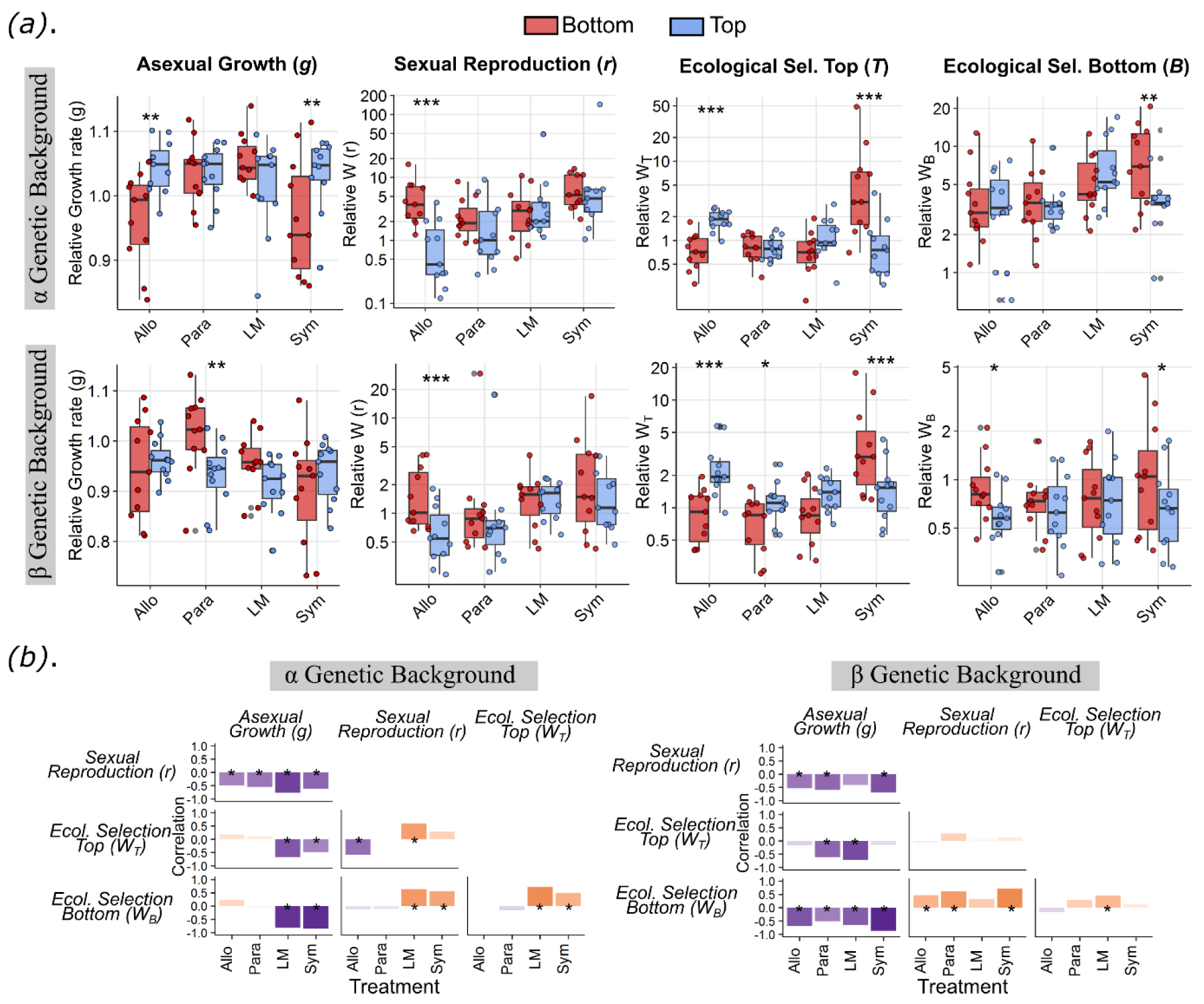
Fitness as a function of ecological selection and gene flow. **a**. Fitness values of each population relative to the ancestral population for different fitness components (growth, sexual reproduction, performance after top or bottom selection) separated by ecological selection regime (top in blue and bottom in red; cf. **Figure 1b**), migration treatment (allopatric, parapatric, local mating, sympatric) and genetic background (α or β). Values > 1 provide evidence for adaptation. Each point represents the median value of 8 technical replicates per population. Note that for local mating and sympatry the entire population experiences top and bottom selection. Blue and red here refers to fitness values of the resulting top and bottom fractions (ecotypes) that experienced two additional cycles of selection in isolation. Statistical significance of Generalized Linear Model is indicated by asterisks (for statistical results see **Supplementary Table 1**; *p < 0.05, **p < 0.01, ***p < 0.001). Boxplots description: center line, median; box limits, upper and lower quartiles; whiskers, 1.5x interquartile range; points, outliers. **b**. Matrix of correlation coefficients of relative fitness values between fitness components per treatment and genetic background (Pearson’s correlation across all 22 evolved populations (allopatry, parapatry) or subpopulations (local mating, sympatry); *p<0.05). The direction and strength of the relationship is indicated by colour (violet: negative; orange: positive) and the height of the bar, respectively.

Populations that experienced intermediate levels of migration (‘parapatry’ and ‘local mating’) showed a different response. Even though these populations showed changes relative to the ancestor, no divergence between selection regimes was observed for any of the four fitness components (**Figure 2a**). Moreover, in these two treatments, fitness values for the populations evolving as pairs were strongly correlated and highly similar (**Supplementary Figure 3, Supplementary Table 1 & 2**). These results support the prediction that intermediate levels of gene flow tend to homogenize population pairs and constrain divergence ^10,33,9^.

In contrast to expectations that higher migration reduces diversification, sympatric populations, which experienced the highest level of migration, showed evidence for divergence when separating each population into two fractions (top and bottom) followed by two continued cycles of disruptive selection without gene flow (see **methods and discussion below**). Similar to divergence between populations under allopatry (i.e. no gene flow), divergence was evident between sympatric top and bottom fractions for three of the fitness components (*g, W*_*T*_, *W*_*B*_). In contrast to allopatry and expected for a high gene flow regime^34^, the bottom fraction had evolved a generalist survival strategy outperforming the top fraction in both the top and bottom environment **(Figure 2a**). Importantly, however, the top selected fraction showed increased fitness only during asexual growth, suggesting the emergence of a polymorphism between an ecological generalist and a growth specialist (see below). The local mating treatment expected to foster divergence ^35,36^ did not result in similar ecotypic sub-functionalisation. Results from the β background were comparable, showing most divergence in allopatry and sympatry, followed by parapatry and least in the local mating treatment (**Figure 2a & Supplementary Figure 3**). In summary, adaptive divergence evolved more readily in the α genetic background and was most pronounced at the extreme end of the migration gradient. In allopatry, it resulted in the emergence of top and bottom specialists, whereas in sympatry we observed a polymorphism between an ecological generalist and a growth specialist.

### Genetic correlations and G-matrix evolution

Consistent with expectations ^25,12^, evolutionary responses of the various fitness components were governed by intrinsic correlations. To investigate the direction, strength and evolutionary stability of these correlations, we constructed standardized variance-covariance matrices (G-matrices) for all four fitness components: asexual growth rate (*g*), reproductive success (*r*) and survival to top (*W*_*T*_) or bottom selection (*W*_*B*_) (**Figure 2b**). Positive parameter values for a given pair of components indicate consistent evolutionary responses across all evolved populations for both fitness components. Negative parameter values indicate an increase in one component accompanied by a decrease in the other component, suggesting a classical trade-off. We observed a pervasive negative relationship between growth rate and sexual reproductive success across all migration treatments and genetic backgrounds (**Figure 2b**) despite large differences in growth rate between ancestral populations (**Supplementary Figure 2)**. Other relationships, such as a negative correlation between survival during top selection and growth, or a positive correlation between survival during bottom selection and sexual reproduction were dependent on the amount of migration (**Figure 2b**). Additionally, correlations between fitness components were contingent on the genetic background. Correlations were more consistent for different migration treatments in the β background possibly due to the lower degree of adaptive divergence observed for these populations. Analysing the covariance in fitness components using Principal Component Analysis with normalized fitness values confirmed these findings (**Supplementary Figure 4**). In summary, phenotypic evolution was strongly governed by intrinsic correlation and potential trade-offs, but correlations between fitness components varied in their stability and were influenced by the level of gene flow and genetic background.

### Genetic correlations and adaptive divergence

Above, we have shown that adaptive divergence was most pronounced in allopatry and sympatry, particularly in the α genetic background. In allopatry, directional ecological selection in a given environment elicited a correlated response in sexual reproduction (α & β background) and growth (α background). The direction of the response defined top or bottom specialists (top: high fitness for growth, low for sexual reproduction; bottom: low for growth, high for sexual reproduction; **Figure 2a**). In contrast, populations exposed to maximal levels of migration (sympatry) experienced both top and bottom selection environments during each cycle and developed a generalist ecological strategy. This generalist strategy showed high performance in both environments as well as in sexual reproduction (**Figure 2a**). However, due to the pervasive trade-off of these components with asexual growth (**Figure 2b**), populations could not increase fitness for all life-history components simultaneously. In populations of the α background, this trade-off was consistently resolved by an intra-population polymorphism: a bottom generalist performing well in both environments and sexual reproduction, and a top specialist with high performance for speed during asexual growth (**Figure 2a**). Overall, these analyses highlight the importance of gene flow on the evolution of genetic correlations and their impact on adaptive divergence.

### Migration and the partitioning of genetic variation

To investigate the genetic basis of adaptive evolutionary change, we inferred population allele frequencies for all 132 evolved populations. At the beginning of the experiment, we identified 107 and 114 genetic variants (SNPs and small indels) representing standing genetic variation in the α and β ancestral genetic background, respectively (**Supplementary Figure 5**). After 53 sexual generations, we counted a total of 1,472 (α background) and 1,318 variants (β background) ranging from 71 to 183 variants per population. Most variants were present at low frequencies (∼80% with maximum frequency < 0.2), and/or limited to single populations (∼62% of all variants) (**Supplementary Figure 6**).

To test for genetic differentiation between top and bottom environments, we first performed Principal Component Analyses (PCA) using allele frequencies of allopatric populations. We observed that genetic variation of allopatric populations was partitioned according to selection regime. Top and bottom populations diverged from the ancestor in opposite directions along the main axis of variation (**Figure 3a, Supplementary Figure 7**, linear model on PCA1 by ecological selection regime, p<1.0 × 10^−3^ for both genetic backgrounds). The clustering by selection regime is best explained by parallel allele frequency shifts of standing genetic variation due to ecological selection. Separation of populations into top and bottom clusters thus provides evidence for allelic separation of shared, ancestral genetic variants by top and bottom ecotypes (further discussed in ‘Genetic architecture of genetic variation’). Consistent with this interpretation measures of genetic divergence, *D*_*xy*_ and *F*_*st*_, were highest between top and bottom ecotypes only when considering standing genetic variation (**Supplementary Figures 8 & 9)**.

**Figure 3.**
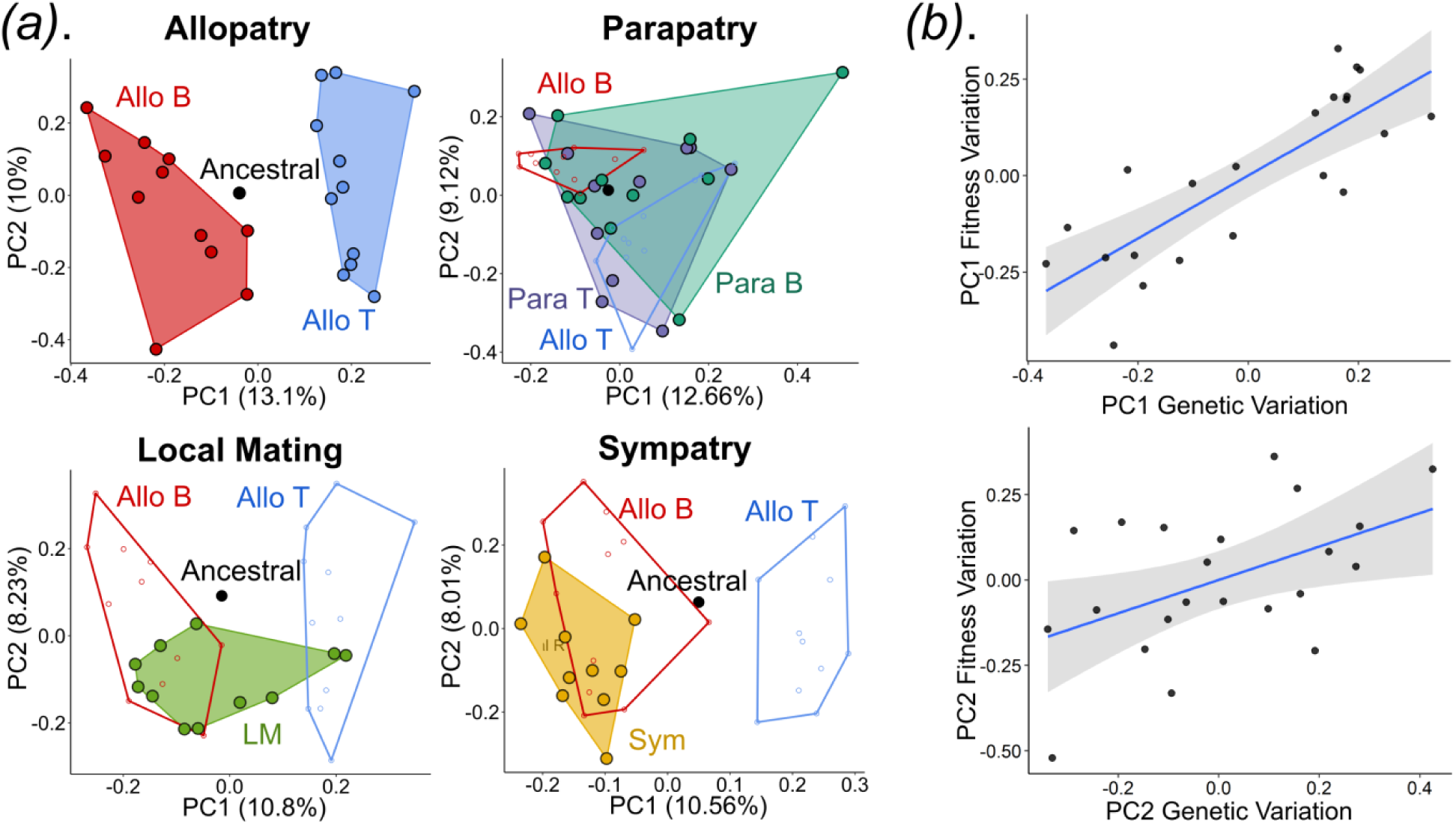
Partitioning of genetic variation and its relationship to fitness. **a**. Genetic variation and population structure relative to the ecological selection regime shown for the α genetic background. Each sub-plot shows the two main axes of variation across populations per treatment always including allopatric top and bottom as well as the ancestral population as reference (blue, red and black, respectively). PCA was performed on all genetic variants. For PCA across all populations combined for each ancestral background see **Supplementary Figure 7. b**. Correlation between the two major PCA axes of genotypic variation (from **Figure 3a**) and variation in fitness (**Supplementary Figure 4**) shown for allopatric α populations. Each dot represents one population.

In contrast, parapatric populations showed no consistent genetic differentiation between connected top and bottom population pairs neither for standing variation (**Figure 3a, Supplementary Figure 7**) nor genome-wide (**Supplementary Figures 8 & 9)**. Instead, pairs of parapatric populations were genetically more similar to each other than to populations from the same selection regime suggesting homogenization by gene flow. Yet, while gene flow inhibited genetic divergence, it did not exclude evolution per se. Genetic variation across independent population pairs was comparable to the genotypic space (PC1 and PC2) of allopatric top and bottom specialists (**Figure 3a, Supplementary Figures 8 & 9**). Local mating populations also spanned a broad range of genetic variation, though clustered primarily with allopatric bottom populations (**Figure 3a**). Populations evolving in sympatry exclusively carried a signature of genetic variation characteristic for allopatric bottom populations (**Figure 3a**, Sym_Allo_B vs. Sym_Allo_T in **Supplementary Figures 8 & 9**).

These findings suggest that a moderate amount of gene flow (parapatry) still allows populations to accumulate genetic variation that is beneficial to both ends of the disruptive selection regime, but precludes adaptive divergence between connected population pairs. With increasing levels of gene flow, however, one ecological condition (here bottom selection regime) appears to dominate the component of standing genetic variation responding to selection. While this pattern is consistent with the observed increase in fitness for bottom selection in the sympatric bottom ecotype, it fails to explain the simultaneous increase for fitness components responding to top selection (sympatry) or the shift in the G-matrix (local mating, sympatry). In sympatry, where phenotypic data suggest coexistence of an ecological generalist bottom ecotype and a growth specialist (**Figure 2a)** this implies an additional role of population-specific novel mutations allowing divergence of these strategies.

In order to test for genetic divergence between ecotypes, we additionally sequenced subpopulations after two rounds of top and bottom selection from sympatric and local mating populations. In the α background, net sequence divergence (*D*_*a*_, ^37^) was positive for all populations indicating genetic differentiation between top and bottom ecotypes. Genetic divergence was higher in sympatry compared to local mating mirroring patterns of phenotypic divergence (**Supplementary Figure 10**). Sympatric divergence was further characterized by genetic variants increasing in frequency during both top and bottom selection, with a significant skew towards top selected variants which was not observed in local mating populations (**Supplementary Figures 11 - 13**). Moreover, selection-induced changes in allele frequencies were less pronounced after top selection suggesting that the pool population may be dominated by variants beneficial to the top ecotype (**Supplementary Figures 11 - 13**). Assuming no systematic bias in effect sizes between variants, these results are consistent with the idea of co-existing strategies in α sympatric populations rather than exclusive dominance of a bottom generalist ecotype. In contrast, for sympatric ecotypes of the β background, net divergence was similarly low as for local mating populations, and allele frequency shifts showed no major contribution of top selected variants (**Supplementary Figure 10 - 13**). In conjunction with the phenotypic data, these results may indicate that these populations are mainly dominated by a bottom generalist ecotype lacking evidence for polymorphism with a growth specialist (**Figure 2a)**. Interestingly, these data suggest that local mating also contains genetically differentiated subpopulations despite clear divergence for the life history traits we measured. In summary, the genetic data suggests different modes of adaptation depending on the degree of gene flow. In allopatry, ecological specialization resulted from parallel allele frequency shifts of standing genetic variation, whereas in sympatry (α background) independent, novel mutations appear to repeatedly induce a polymorphism of strategies.

### Genetic architecture of adaptive variation

As expected from the large experimental population sizes, the genetic composition of evolved populations was found to be governed by ecological selection, and not genetic drift. In allopatric populations, the importance of selection for driving the correlation between traits was illustrated by the fact that the major axes of genetic variation (**Figure 3a**) and variation in fitness components (**Supplementary Figure 4**) were strongly correlated (**Figure 3b**, PC1: R_adj._^2^ = 0.69, p < 0.0001, PC2: R_adj._^2^= 0.16, p = 0.021). In the presence of gene flow (parapatry), the correlation was reduced and only significant for the main axis of variation (PC1: R_adj._^2^ = 0.15, p = 0.037; PC2: R_adj._^2^ = 0.04, p = 0.31). In β populations, where the divergence between allopatric top and bottom populations was weaker, the correlation was not statistically supported (PC1: p = 0.1, R_adj._^2^ = 0.07; PC2: p = 0.7, R_adj._^2^ = 0.04). Individual based simulations mirroring the allopatric experimental setup further supported that the observed increase in allele frequency of novel mutations was the result of ecological selection rather than drift (**Supplementary Figures 14 & 15**). Moreover, mutations with low predicted functional effects (synonymous sites, non-coding regions) segregated at low frequencies, whereas mutations with moderate (missense variant, codon loss/gain) and strong predicted effects (frame shift, stop gain, start loss) increased to significantly higher frequencies in all populations (**Supplementary Figure 16**). This disproportionately strong increase in frequency of mutations with strong predicted effects was pervasive across all levels of migration (**Supplementary Figure 17**).

Next, we quantified the degree of parallelism in allele frequency shifts. 34 and 50 genetic variants corresponding to 32 % and 44% of all standing genetic variation from the α and β background, respectively, showed consistent differences in the direction of allele frequency changes between allopatric top and bottom populations (**Supplementary Figure 18**). However, only five (α) and three (β) genetic variants reached frequencies above 0.9 for at least two populations in one ecological regime while going extinct in the opposite regime for most populations. These same variants contributed the main loadings on Principal Component 1 of the overall genetic variation across treatments suggesting a major role in the evolutionary response irrespective of the amount of migration (**Supplementary Figure 19)**. Evolutionary parallelism was not restricted to single sites of standing genetic variation, but was also observed for novel mutations at the gene level. 140 genes significantly enriched in GO-terms for cell-cell adhesion (flocculation and agglutination), polysaccharide catabolic process and cell cycle regulation were hit by multiple novel mutations (up to 187 variants per gene, **Supplementary Figure 20 and Supplementary Table 3**). Cell adhesion traits can increase cluster formation, which might increase settling speed ^38^ or improve sexual reproduction. Most of the genetic variants, however, did not reach fixation, which may be attributed to genetic redundancy, size effect distribution, negative epistasis, antagonistic pleiotropy or balancing selection ^39,40^.

A single genetic variant that is beneficial in one environment can be beneficial, neutral (conditional neutrality) ^41,23,42^ or deleterious (antagonistic pleiotropy) ^43,44^ in another environment. Under conditions of gene flow, allele frequency differences between populations are more likely to be maintained under antagonistic pleiotropy ^45^. Additionally, if several loci are subject to antagonistic pleiotropy, linkage disequilibrium can arise even in the absence of epistasis and form the basis for reproductive isolation ^46,47^. Even though our pool-seq data does not provide haplotype information, we found evidence for multilocus antagonistic pleiotropy of closely linked loci (**Supplementary Figure 18**), which appears to commonly arise under divergent selection ^6,42,44^. The strongest and most consistent allelic differentiation caused by the ecological selection regime was found for a neighbouring pair of mutations on chromosome II (22kb distance) in the genes *rep2* (variant II:1718756_A; early stop C178*) and *byr2* (variant II:1741521_T; amino acid substitution I259N) in the α genetic background. Both genes are involved in cell cycle regulation, either during mitotic (*rep2*) or meiotic (*byr2*) reproduction. The derived allele for *rep2* had consistently elevated frequencies in allopatric bottom populations relative to the ancestral α population (p_rep2_=0.5), but reduced frequencies in top populations (p_byr2_=0.2). The derived *byr2* allele showed the opposite pattern. Not a single replicate population showed simultaneous positive selection of both derived alleles (grey area in **Figure 4a**). With the exception of two local mating populations this held true across migration treatments (**Supplementary Figure 21)**. Moreover, we observed no incidence where the sum of the derived allele frequencies of both loci would exceed a value of 1, which would provide unequivocal evidence for coupling of derived mutations in one haplotype (diagonal line in **Figure 4**). Deterministic simulations further supported opposite directional selection of both loci, as opposed to a scenario of selection on one locus and hitchhiking of a neutral linked variant (see **Methods** and **Supplementary Figures 22**). Overall, these results provide evidence for multilocus antagonistic pleiotropy of two derived mutations being favoured in opposite environments (**Figures 4b and 4c**).

**Figure 4.**
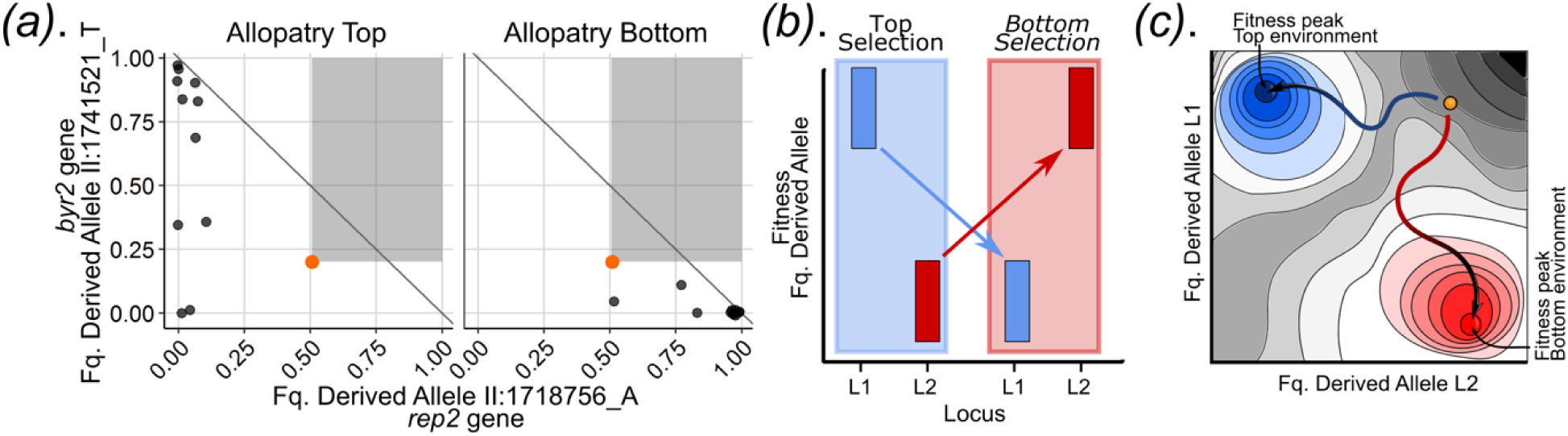
Candidate genetic variants under disruptive selection in α populations. **a**. Example of a pair of variants with evidence for antagonistic pleiotropy. Each point shows the final allele frequency for the two loci in top or bottom populations from the allopatric treatment (frequencies for other treatments are shown in **Supplementary Figure 21**). Ancestral allele frequencies are shown in orange. Genetic variants are labelled with chromosome number, base position and alternative allele relative to the reference genome. The *grey area* indicates allele frequency combinations that would arise under a scenario of positive selection for both derived alleles; allele frequency combinations above the *diagonal line* provide unequivocal evidence for coupling of both derived mutations on a single haplotype. **b**. Predicted allele distribution for a pair of genetic variants under antagonistic pleiotropy. The derived allele for locus 1 (L1) is beneficial under environment 1 (top selection environment) but deleterious in environment 2 (bottom selection environment). The opposite occurs for locus 2 (L2). **c**. Representation of a fitness landscape where a population (orange circle) can adapt to the top or bottom environment (blue or red arrow, respectively) by increasing the frequency of the derived allele at locus 1 while decreasing the frequency for the derived allele at locus 2.

In the β background, a pair of loci with comparable dynamics was found within a single gene, *msa1*, with the variants I:2319886_T (W106*; high frequency in bottom populations;) and I:2319922_T (W118*; high frequency in top populations) each introducing an early stop codon (**Supplementary Figures 23 & 24**). Similar to *byr2*, wild type *msa1* suppresses sporulation. Evidence of antagonistic pleiotropy is rare in natural populations, but is expected to favour local adaptation and reproductive isolation ^45^. The occurrence of several tightly linked loci showing antagonistic pleiotropy (e.g. locked in an inversion ^48^) is of particular interest in the context of speciation, as the joint effects of multiple loci increase the potential for coupling of these effects, inducing reproductive isolation ^47,49^.

## Summary and Conclusion

This study provides experimentally controlled, empirical insight into the effect of migration on adaptive divergence. Parallel divergence was readily achieved in isolation as expected under opposing directional selection ^11^, mostly from standing genetic variation. Intermediate levels of homogenizing gene flow reduced divergence, but the occupied trait space and genetic variation between population pairs encompassed the full range of locally adapted allopatric populations ^10,50^. Contrary to theoretical expectations and previous empirical findings ^19,51,52^, this included the local mating treatment where sexual reproduction was matched by environment expected to act as a source of premating isolation. Moreover, in contrast to many studies in natural systems, we also did not observe intermediate divergence at intermediate levels of migration (isolation-by-distance and isolation-by-ecology relationships) ^8,17,22,53,54^. This is likely owing to the combination of short divergence time and rather high levels of migration (even in parapatry) and the near-absence of genetic drift in our experimental setup. Complementary experiments^23^ or sampling of natural populations across a finer-scale of intermediate migration levels ^18,55,56^ are thus highly encouraged.

Contrary to the parapatric and local mating treatment, adaptive divergence was apparent under maximal levels of migration. This is a puzzling observation running counter to the general expectation of an inverse relationship between gene flow and divergence ^14^. Under conditions of high gene flow adaptive divergence implies emergence of (quasi-)stable co-existence of distinct strategies allowing exploitation of different niche space ^23^. Independent accumulation of mutations promoting further divergence then constitutes the basis, though no guarantee, for ecological speciation^19^. The general expectation, however, is that high levels of gene flow may promote the evolution of an ecological generalist gradually taking over the population without promoting population divergence ^57–59,34,60^. Under the latter scenario, the polymorphism observed in the sympatric treatment might be the result of directional selection for a generalist strategy which has not yet reached fixation during the course of 53 sexual generations. Results from several replicates of β sympatric populations are possibly consistent with this scenario. In the vast majority of replicate populations from the α background, however, several lines of evidence support the evolution of true adaptive divergence in sympatry where the pervasive negative correlation and the evolvability of genetic correlations appear to be key. In addition to the evolution of an ecological generalist with high survival in both environments, the strong trade-off with asexual growth additionally promoted the emergence of a second strategy specializing on performance during asexual growth (**Figure 2**). As a consequence, multivariate phenotypic divergence integrating across life history components reached levels comparable to allopatric populations and exceeded those of parapatry and local mating. At the genetic level, a consistent increase of functional genetic variation (D_a_) between ecotypes across nearly all α sympatric populations, but few of the local mating populations sharing the same ancestor, lends further support to the parallel existence of distinct adaptive types independently accumulating mutations. At the current stage, we are ignorant about the long-term stability of these types. However, the fact that they repeatedly evolved and could be observed in nearly all sympatric populations (including several populations of the β background) speaks against a transitory sweep pushing a single, generalist strategy to dominate. Moreover, the relatively large shift in allele frequencies observed in sympatric populations after two rounds of directional selection (**Supplementary Figures 13**) is difficult to reconcile with a transitory sweep and rather suggests maintenance of polymorphism by divergent selection.

Stable co-existence of both strategies may be achieved by two non-exclusive mechanisms: (i) by assortative mating facilitated by the evolution of self-compatibility (observed in our experimental setup ^61^) or temporal asynchrony in sporulation as anecdotally observed for a subset of populations; (ii) by antagonistic pleiotropy of large effect genes or strong negative epistasis of alleles coding for different life history components ^62–65^. Regardless of the precise mechanism, this result overall exemplifies the importance of genetic correlations for inhibiting or enabling adaptive divergence with gene flow, an aspect that may deserve more attention both from a theoretical viewpoint ^12,13^, as well as in empirical studies of natural systems ^20,25^.

Consistent with contributions of several fitness components to adaptive divergence the underlying genetic basis was polygenic ^25^. In line with theoretical predictions ^66,67^ and existing empirical studies ^39^, effect sizes from standing genetic variation were skewed with only a small, repeated fraction of genes showing large effects on population differentiation. This was most pronounced in allopatry, where parallel allele frequency shifts of standing genetic variation governed adaptation. In sympatry, populations were near-exclusively characterized by standing genetic variation of allopatric bottom populations. We speculate that this similarity may help explain why diversification more readily occurred in the sympatric migration treatment. Assuming initially stronger selection in the bottom environment the high degree of gene flow might have moved much of the population towards a single (bottom) strategy increasing local competition and opening the opportunity for a top specialist to invade ^16^. In the intermediate treatments, a generalist strategy might have been maintained inhibiting the evolution of a specialist strategy. Divergence in sympatry requires the emergence of novel mutations conferring the necessary variation promoting concurrent evolution of an ecological generalist and a growth specialist. This is consistent with our observation of a multitude of unique, novel mutations which were, however, concentrated in specific genes and functional pathways. As a consequence, evolutionary trajectories may be partly predictable, not only at the phenotypic level, but also at the level of the underlying genes ^68,69^. Future work unravelling the genotype-fitness map will be necessary to understand the genetic architecture of the opposing adaptive strategies and their trade-offs.

Concluding, this evolve-and-resequence experiment demonstrates that divergent selection readily promotes adaptive divergence of ecological specialists in the absence of migration and facilitates the evolution of ecological generalists under conditions of gene flow. Importantly, it further provides evidence that adaptive divergence is also possible, if not favoured, under maximal levels of migration, whereby the evolution of genetic correlations of fitness components appears to play a vital role. The genetic basis of divergence was conferred by a large number of genes exploiting both standing genetic variation in major effect genes with evidence for antagonistic pleiotropy and novel mutations enriched in certain genes and metabolic pathways. These findings contribute to our understanding of the fundamental processes governing adaptation and have potential implications for speciation research, pathogen evolution, pest control or conservation biology.

## Methods

### Ancestral populations

The preparation of the α and β ancestral populations started with four isogenic strains (parental strains: P1, P2, P5, P6) derived from the Leupold’s 968 accession ^70^. These four parental strains differed in 14-22 genetic variants (**Supplementary figure 25**) including mutations in the *ade6* gene used as colour marker (*ade6-M216* allele in P1 and P5, and the *ade6-M210* allele in P2 and P6), and the homothallic mating locus with configuration *h*^*-S*^ (in P2 and P5) or *h*^*S*^ ((in P1 and P6 ^71^). These strains are obligatory outcrossing (heterothallic), although along the experiment we observed the emergence of homothallic phenotypes (mating type switching), ^61^. In general, unless specified, asexual growth was performed in standard liquid Edinburgh Minimal Medium (EMM; Per liter: Potassium Hydrogen Phthalate 3.0 g, Na HPO_4_·2H_2_O 2.76 g, NH_4_Cl 5.0 g, D-glucose 20 g, MgCl_2_·6H_2_O 1.05 g, CaCl_2_·2H2O 14.7 mg, KCl 1 g, Na_2_SO_4_ 40 mg, Vitamin Stock ×1000 1.0 ml, Mineral Stock ×10,000 0.1 ml ^72^). Sexual reproduction took place on 2% agar solid Pombe Minimal Glutamate medium (PMG corresponding to EMM medium substituting ammonium chloride by 5 g l^− 1^ glutamic acid). Ecological selection was conducted in Selection Medium (SM; as PMG with glutamic acid reduced to 0.8 g l^-1^). In all cases, media was supplemented with 100 mg l^− 1^ adenine.

Preparing the ancestral populations for the experiment involved two phases. In the first phase (*phase I*), settling speed at stationary phase was used as ecological contrast resulting in disruptive selection on a complex trait involving growth rate, cell size and cell morphology ^38^ (**Supplementary Figure 15**).^37^ (**Supplementary Figure 26**). The aim of *phase I* was to induce genetic variation relevant to ecological specialization for fast or slow settling rate (bottom and top selection regime, respectively) in independent, asexually reproducing populations. In total, we maintained 24 populations (6 populations per parental strain), 12 for fast settling (bottom selection) and 12 for slow settling (top selection). The experiment was performed in cycles of asexual growth in 5 ml of EMM at 32°C shaking at 250 rpm for two days, followed by a selection step where 1% of the cells were transferred to fresh medium (around 5 million cells were transferred). For bottom selection, 1 ml of saturated media were placed on the top of a column with 10 ml of SM. The column was centrifuged for 45 seconds at 100 g, and 300μl from the bottom fraction of the column were collected. For top selection, the saturated medium was diluted to a final volume of 15ml with water, and centrifuged for 2 minutes and 45 sec at 100 g, after which 500 μl of the surface liquid was collected. Collected bottom and top fractions were then placed in 5 ml of fresh EMM media for each population to start a new cycle. In total, we conducted 50 and 62 selection cycles for bottom and top selection, respectively, corresponding to approximately 430 and 530 asexual generations. At the end of this phase, each of the 24 evolved populations was diluted and plated in solid EMM with 2% agar. Plates were grown for three days and one single colony was isolated from each population (one isogenic strain per evolved population) to initiate the second phase (*phase II*).

Since the *phase I* was run only using asexual cycles, during *phase II*, the aim was to add sexual reproduction to the cycles and increase reproductive efficiency, while maintaining variability in the ecological selection regime (top and bottom selection) produced in *phase I. Phase II* was started in duplicate maintaining the identity of the colour marker using the strains derived from evolved populations from *phase I* with parental P2 and P6 ancestry (*ade6-M210* allele, or *α* genetic background) or from parent P1 and P5 (*ade6-M216* allele, or *β* genetic background). Each of the 24 evolved strains were grown to saturation in EMM, and within the α and β genetic background the six Plus mating types were each mixed in equal proportion to each of the six Minus mating types for all possible combinations, and thereafter transferred to mating plates. After three days of mating on solid PMG, 1% of the cells were harvested from each cross, sexually produced offspring (ascospores) were isolated by killing all non-mated cells using Glusulase (0.5% v/v overnight; PerkinElmer) followed by a 30% ethanol treatment for 30 minutes. Spores were recovered and incubated in 5ml of EMM at 32°C for two days. A second round of mating was performed, now mixing all offspring per genetic background from the first round resulting in two population pools (α and β). For each pool ten independent replicate populations were propagated for 20 cycles of disruptive selection described above with the addition of sexual reproduction after the selection step (**Supplementary Figure 27**). Sexual reproduction was introduced in two ways: in half of the populations (five each α and β) cells were mixed prior to sexual reproduction, and in the remaining populations mating was performed independently in each of the selection fractions (bottom or top) where after spores were mixed in equal proportions. In both treatments spores were harvested to remove un-mated cells by glusulase and ethanol treatment as described above and were used to start the new cycle. *Phase II* was run for 20 cycles, each lasting six days. After 20 sexual generations, the 10 evolved populations from each ancestral α or β population were mixed to produce two independent populations with different genetic background and different composition of standing variation. These populations were used to start the experiment forming the basis of this study. They are referred to (α and β) ancestral populations.

### Experimental evolution: divergent selection with migration

The evolutionary experiment was run in duplicate using both ancestral population (α or β populations from end of *phase II*). For each background, we ran 66 replicate populations corresponding to four treatments varying in the level of migration (see below). All steps of the experiment were performed in 96-well plates, either 1.2ml deep-well plates for asexual growth and ecological selection, or flat bottom 360 μl microtiter plates for sexual reproduction. Similar to preparation *phase II* the experiment was run in six day cycles, each including growth, ecological selection and sexual reproduction. Experimental conditions were modified to accommodate larger numbers of replicates and introduce four levels of migration. Populations were grown asexually for two days in 300 μl of EMM per population followed by ecological selection. For bottom selection, 50 μl of cells were placed on the top of a column with 750 μl of SM in a 96 deep-well plate. After 9 minutes, 25 μl of the bottom fraction was collected (corresponding to 0.5 % of cells, or around 150,000 cells). For top selection, 100 μl of cells were placed on the top of a column with 550 μl of SM in a 96-well plate. This plate was centrifuged at 100 rcf (705 rpm) for two minutes and 45 seconds, and 25 μl were collected from the surface (again corresponding to 0.5 % of cells, or around 150,000 cells). The subsequent step of sexual reproduction was performed in microtiter plates with 150 μl of PMG per well. After three days, 25% of the sexually produced offspring (ascospores) were harvested by killing all non-mated cells using Glusulase (digestive enzyme mixture) in the incubator at 27.5 °C overnight, followed by 30% ethanol treatment for 30 minutes. These spores were used to start the next cycle.

For each background, we modified the amount of migration between ecologically selected populations (top and bottom fraction after selection) in four treatments (**Figure 1 and Supplementary Figure 1**). i) In the *allopatric* treatment, half of the populations were subjected to bottom selection, and half to top selection. Sexual reproduction was restricted to within each population. ii) In the *parapatric* treatment, replicates were divided into non-independent population pairs experiencing opposite ecological selection (top or bottom selection). After selection, 5% of the selected cells were reciprocally transferred between populations of each pair. Sexual reproduction occurred independently in each population. iii) In the *local mating* treatment, independent populations were grown asexually and experienced disruptive selection for both, top and bottom selection. Sexual reproduction occurred in each resulting fraction independently. The spores produced from top and bottom mating plates were then mixed and transferred together for asexual growth. iv) In the s*ympatric* treatment, independent populations were grown asexually and experienced disruptive selection for top and bottom selection. Prior to sexual reproduction the two fractions were fully mixed, transferred to a mating plate and the resulting spores were used again for asexual growth.

Each treatment contained 22 replicates of evolving populations (sympatry and local mating treatment) or pairs of populations (allopatry and parapatry) giving a total of 66 replicate populations for each ancestral background, or 132 altogether. In order to maintain the same population mutation rate (*N*_*e*_*μ*) in all treatments, population size was matched during asexual growth performed in two independent wells per population for the sympatric and local mating treatments, which were then mixed after growth before selection. Additionally, the sexual reproduction step for the sympatric treatment was performed in two wells per population. In order to control for cross contamination between populations, we included two empty wells per treatment, with media, but without cells. The experiment was run in total for 53 cycles (53 cycles of sexual reproduction and around 700 asexual generations).

During the entire experiment, populations were stored every three cycles after asexual growth in YES medium with 15 % glycerol, and were cryopreserved at -80°C.

### Fitness measurements

We measured fitness of the evolved populations relative to the ancestral populations (α and β ancestral populations after the two preparation phases) for four fitness components: asexual growth, sexual reproduction and survival of ecological selection for top or bottom. Additionally, in order to identify the potential for differentiation within populations (subpopulations - population structure), each population was subjected to two experimental cycles removing migration (as in allopatric treatment). This resulted in a top and bottom subpopulation. To ensure comparability across treatments the two additional cycles were performed prior to fitness measurement in all four treatments. To quantify relative fitness, we performed a competition assay between all evolved populations and ancestral populations (test populations) using a fluorescent isogenic strain as intermediary reference. The reference fluorescent strain was derived from the lab strain Leupold’s 968 h^90 70^ containing an introduced mCherry-marker (strain EBC47 described in^61^). Each test population and reference fluorescent strain was first grown independently to saturation for two days. Subsequently, 250 μl of the test population were mixed with 100 μl of the reference (mix before growth - BG). 10 μl of the mix were then diluted with 100 μl of water and the frequency of fluorescent (reference strain) vs. no-fluorescent cells (from the test population) were measured using a flow cytometer (BD LSR Fortessa, at the Core Facility Flow Cytometry, LMU) (baseline frequency before growth – *f*_BG_). A fraction of 5 μl (BG) was then mixed with 500 μl of SM (selection media) of which 30 μl were transferred to 100 μl of EMM. After 24 h of asexual growth, the change in frequency was measured using the flow cytometer (frequency after growth – *f*_AG_) providing an estimate of growth rate differences between the test population and the reference. Another fraction of 50 μl (BG) was subjected to one cycle of bottom selection. Given the reduced number of cells for measurements after selection, the selected fraction was mixed with 100 μl of EMM and grown for 24 hours. After growth, the change in frequency was measured using the flow cytometer (frequency after bottom selection plus growth – *f*_BSG_). A third fraction of 100 μl (BG) was placed on the top of a top selection column, and a cycle of top selection was performed. The selected fraction was grown and measured as for bottom selection described above (frequency after top selection and growth – *f*_TSG_). To measure sexual reproductive success, 20 μl of the reference strain and 80 μl of the test population were mixed, and the fluorescent and non-fluorescent proportion of this mix was measured in the flow cytometer (frequency before mating - *f*_BM_). To reproduce the evolutionary environment, 10 μl of the mix was diluted to 1,000 μl of SM of which 25 μl was transferred to a PMG mating plate. After three days of sexual reproduction, spores were harvested as in the experiment and transferred to 100 μl of EMM for asexual growth. After 24 hours, samples were measured in the flow cytometer (frequency after mating and growth – *f*_AMG_). Eight technical replicates were performed for each fitness component measurement and population. Raw data was converted using flowCore 1.11.20 (Ellis et al. 2009) and analysed in R. Debris was filtered by gating in FSC width and height and a cut-off in the mCherry signal was used to define reference and focal populations (see **Supplementary Figure 28** for representative example).

All fitness components were measured relative to the reference fluorescent strain. Due to technical limitations, measurements required a growth step after selection and after sexual reproduction. To compensate for these steps, we used the calculations described below, to obtain fitness estimates of the evolved populations relative to the (α or β) ancestral population of the experiment (adaptation). First, we estimated the number of asexual generations for the reference fluorescent strain in 24 hours to be around 6. Given an initial frequency of cells before growth (*f*_BG_) for both evolved population and reference strain, as well as the frequency after growth (*f*_AG_), we inferred the number of reference cells after 24 hours of growth as:

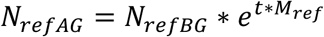

Where *t* = 6 and *M* _*ref*_ = 1 (Malthusian param eter of the r eference strain was s et to 1). The number of cells of the evolved populations is:

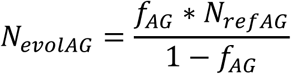

Then the number of evolved cells before and after growth were used to calculate a Malthusian growth parameter for the evolved population as:

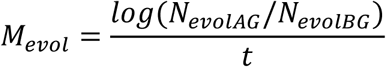

The same calculation was used to estimate a Malthusian parameter per cell division for the ancestral populations (M_ancestral_). The relative fitness for growth (g) then is:

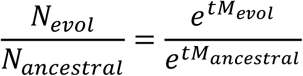

The relative fitness after ecological selection (top or bottoms selection), was calculated in the same way in both cases. We used the calculated M_evol_ parameter to differentiate the change in frequency by selection (frequency after top or bottom selection – *f*_TS_ or *f*_BS_) from the change from selection plus growth (*f*_TSG_ and *f*_BSG_ described above). For that, we calculated the number of reference cells remaining after top and bottom selection alone from saturated medium (Expected_pS_ref_), and used it to calculate the number of reference cells after selection in the mix:

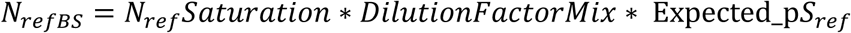

As the fraction after selection was measured after a step of asexual growth of 24 hours, N_refBS_ was used to calculate the number of reference cells after selection and growth (N_refBSG_ and N_refTSG_) with *M*_*ref*_ = 1 as:

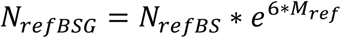

As before, using initial densities and the frequency before selection we calculated the number of evolved cells before selection (N_evolBG_) as:

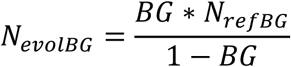

And the number of cells after selection without growth as:

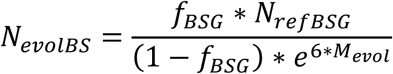

The relative fitness for ecological selection (W) was then calculated as:

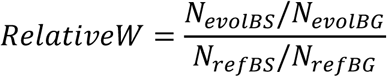

The fitness relative to the reference was calculated for evolved and ancestral populations, and the final relative fitness with respect to ancestral populations was calculated as the ratio between them.

Relative fitness of sexual reproduction was estimated similarly. As the measurements of sexual reproduction efficiency were inferred after a period of 24 hours of asexual growth (AMG), frequencies after mating were corrected using *M*_*evol*_ (frequency after mating without growth – *f*_AM_). Then values of sexual reproduction efficiency of the test population relative to the reference was calculated as:

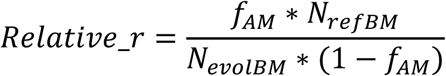

Note that the measure *r* is an aggregate of mating, sporulation and germination efficiency. We assume that the fluorescent marker in the reference strain follows Mendelian inheritance and that the effect of mating between evolved and reference cells is on average equal for the offspring with and without the fluorescent marker.

Estimates of relative growth rate (g), response to selection for top or bottom (W_T_ or W_B_, respectively), and the efficiency of sexual reproduction (r) constitute fitness components that were compared between migration treatments and selection regimes. For each fitness component (ratio data) a log transformation was performed to obtain a normal distribution to which a generalised linear model was fitted using treatment and selection regime as fixed variables and population and technical replicate as random factors. To correct for occasional outliers due to experimental error, for each population, the replicate that deviated most from the median was discarded. In this model, treatments were contrasted to allopatric populations (**Supplementary Table 1 & 2**). In addition, we performed Principal Component Analyses (PCA): i) including all treatments and ii) separately per treatment, using z-score normalized values for the log transformed fitness values per fitness component and visualised using the R package *factoextra* v3.4.4 ^73^; PCAs were conducted for each ancestral genetic background separately, using all variants (**Fig. 3**) or only ancestral variation (**Supplementary Figure 7**). We further calculated correlation matrices of all four fitness components for each treatment and genetic background using standardized z-transformed fitness values. Calculations included all 22 populations (allopatry, parapatry) or subpopulations (local mating, sympatry). **Supplementary Figure 29** shows the correlation for allopatry and sympatry in the α background. The sign of the slope of the lines between points – connecting the pairs of subpopulations in sympatry derived from the same population – indicate if the correlation observed among populations is maintained within populations.

### Genomic data generation and pre-processing

Genetic analyses were performed in all populations obtained during the preparation of the ancestral populations (*phase I* and *II*), the two ancestral populations (α, β), all evolved populations, and subtractions from the local mating and sympatry treatments after two cycles of selection without migration. Specifically, genomic DNA was extracted from the following populations/strains: the four parental strains (P1, P2, P5 and P6), the 24 evolved strains of *phase I*, the two ancestral populations starting *phase II*, the two ancestral populations starting the experiment (α and β ancestors), the 132 evolved populations at the end of the experiment, and 44 samples corresponding to top and bottom fractions from local mating and sympatric populations. Genomic DNA was extracted using Zymo Research Quick-DNA(tm) Fungal/Bacterial 96 Kits according to the manufacturers’ instructions. Library preparation and Illumina HiSeqX, paired-end 150 bp read length, v2.5 sequencing chemistry, was performed at the SNP&SEQ platform of the SciLifeLab at Uppsala University. Libraries were prepared from 1 μg DNA using the TruSeq PCRfree DNA sample preparation kit, targeting an insert size of 350 bp. For sequencing, 48 libraries with barcodes were pooled per lane randomized across treatments. Samples were sequenced to sequence coverage over above 200 x (**Supplementary Figure 30**). Raw sequencing data is available at the National Center for Biotechnology Information (NCBI) under Bioproject ID PRJNA604890.

Adaptors were removed from raw reads using *cutadapt 1*.*3* ^74^, read-pairs were filtered and trimmed by quality using *trimmomatic 0*.*32* ^75^ and *FastQC 0*.*11*.*5*. Filtered reads were then mapped to the reference genome (ASM294v2 ^76^) using *BWA 0*.*7*.*15*^77^. Local realignment was performed using *GATK 3*.*3*.*0* ^78^ and the Picard toolkit *picard 1*.*92* (https://broadinstitute.github.io/picard/). Genetic variants and frequencies (both SNPs and small indels) were inferred using the package *VarScan 2*.*3*.*7* ^79^, with minimum mapping quality of 30 and a threshold p-value of 1×10^−4^. Variants were filtered to exclude 1) variants with more than 90% of the reads supported by a single strand; 2) variants with coverage lower than 10% of genome-wide average or higher than 1.3 times the estimated maximum coverage as suggested by Heng^80^; 3) variants falling into repetitive regions identified using *RepeatMasker 4*.*0*.*7* (http://repeatmasker.org). Genetic variants were annotated relative to the reference genome using *SnpEff 4*.*3*^81^. Based on the annotation for effect on the closest genomic region, genetic variants were classified according to the predicted size effect into four, mutually exclusive categories as defined in *SnpEff 4*.*3*: i) *modifier*, including non-coding transcript exon variant, intragenic region, intron variant, 5’ UTR variants and 3’ UTR variants; ii) *low*, including synonymous, splice region variants, splice region variant and 5’ UTR premature start codon gain variant; iii) *moderate*, including missense variants, disruptive in-frame deletion and insertion, and conservative in-frame insertion; and iv) *high*, including stop-gain, start-loss and frame-shift variant. Coverage values per base were calculated using *SAMtools* 1.9 ^77^. Gene function and phenotype effect for variants that showed strong divergent selection or were hit multiple times independently were checked at pombase.org ^82^. Gene ontology analysis for enrichment was performed using Fisher’s exact test for the annotations of biological process from pombase.org, using false discovery rate as correction.

### Decomposition of genetic and fitness variation

Allele frequencies were used to calculate population genetic parameters including average number of pairwise differences between populations (*D*_*xy*_^83^) and the expected genetic variance within populations relative to the total expected genetic variance (*F*_*st*_^84^) using custom scripts. Population allele frequencies were further used to perform PCAs, which were visualised using the R package *factoextra* v3.4.4. Analyses were conducted for all populations per genetic background (α or β; **Supplementary figure 7**) and by treatment always including the allopatric populations as point of reference (**Figure 3**). This analysis was performed for all mutations (**Figure 3**), or only for variants present in the α or β ancestral populations (standing variation; **Supplementary figure 7**).

We then explored the relationship between genetic variation and variation in fitness components. We extracted the two major axes of variation (PC1 and PC2) for genetic variation (**Figure 3** and **Supplementary Figure 7**) and for variation in the log normalized relative fitness values across all four fitness components (**Supplementary Figure 4**) and investigated the association using a linear model using the *stats v3*.*6*.*0* package. The analysis was done independently for each genetic background and treatment excluding sympatry and the local mating treatments due to the lack of correspondence between sequencing data (from population pools) and fitness estimates (from selected fractions within population – subpopulations). In the case of parapatric populations, the fitted model additionally included ecological selection regime as fixed variable (top and bottom selection regime).

We then evaluated the potential for genetic divergence within populations with high migration (local mating and sympatry). First, allele frequencies were used to calculate ancestral divergence (D_a_: difference between D_xy_ and mean π) between the top and bottom fraction per population (**Supplementary Figure 10**) using custom scripts. Then we compared allele frequency changes between whole pool samples and their respective top and bottom fractions (see examples in **Supplementary Figure 11**). For each population we counted the proportion and number of genetic variants with allele frequency change higher than 0.2 in the comparisons: top – bottom fraction, pool – top, and pool – bottom (**Supplementary Figure 12**). The change of frequency of 0.2 since allele frequency changes higher than 0.15 are not expected for neutral genetic variation (see individual based simulation below), but a lower threshold of 0.1 gave similar qualitative results. Dominant fractions were compared between treatments (local mating and sympatry) and genetic backgrounds (**Supplementary Figure 13**). Significance of the difference between groups were tested using a quasibinomial model in a nested generalised lineal model with treatment and fraction as fixed variables and population as random factors.

### Individual based forward simulations

In order to identify the expected allele frequency distribution of neutral genetic variants and the effect of physical linkage we performed individual based forward simulations using *SLiM 3*.*2*.*1* ^85^. We contrasted simulations including only neutral variants to simulations including both neutral and selected variants We parameterized the simulations with estimates from the literature including a mutation rate of 2·10^−10^ site^-1^ generation^-^1 86^^, an average recombination rate of 1·10^6^ site^-1^ generation^-^1 87^^, and a cloning rate of 0.90 generation^-1^ (equivalent to around 1 sexual cycle every 18 asexual generations). We simulated genetic variation for one chromosome of 1 Mb in size for allopatric populations in cycles following the setup of the experiment (**Figure 1**). 3·10^5^ haploid individuals (cells) grew asexually to a saturation point of 3·10^7^ individuals. Growth was followed by a selection step reducing population size to 3·10^5^, subsequently undergoing a cycle of sexual reproduction with an outcrossing rate of 0.90 generation^-1^. The resulting offspring re-started the cycle. Simulations were run for 800 asexual generations (corresponding approximately to 700 asexual generations during 53 experimental cycles), 1,000 generations (mimicking *phase II* + experiment) and 2,000 generations to explore longer-term evolutionary dynamics. In order to reduce the computational effort, all parameters were scaled relative to an effective population size *N*_*e*_ of 3·10^3^ as suggested in the *SLiM* manual. Simulations were run first including only neutral variants (**Supplementary Figure 14**) and then adding selected variants (**Supplementary Figure 15**). For selected variants, we included a range of parameters specifying: 1) the proportion of emerging selected variants relative to neutral variants (from 100 to 10000 neutral variants per selected variant) and 2) selection coefficients which were sampled from an exponential distribution with varying mean (from 0.01 to 0.1). For each parameter combination, we ran 100 replicate simulations and report the mean tabulated number of genetic variants per allele frequency across simulations. Neutral variants alone did not reach allele frequencies higher than around 0.3 after 2000 generations, and only reached 0.12 in 800 generations (**Supplementary Figure 14**). In the presence of linked selected variants, mean allele frequencies of neutral variants increased, but only under conditions of high selection coefficients and a low occurrence of selected relative to neutral variants (**Supplementary Figure 15**).

### Allelic differentiation by ecological contrast

To test for the difference in allele frequency per variant between ecologically contrasting conditions (allopatric top and bottom regime) we used a logistic regression with binomial and quasibinomial error structure taking over-dispersion into account. In general, the quasibinomial model is not appropriate for cases when variants were only found in some populations (new mutations in the last phase of the experiment), even when derived allele frequencies clearly differed between top and bottom selected populations where present. For example, variants found in high frequency in ∼5 populations, but not present in all bottom populations, where not found to be significantly different in the quasibinomial model. For variants present in all population (standing genetic variation), the quasibinomial model appeared more appropriate.

### Forward simulations for linkage disequilibrium between potentially adaptive loci

This analysis was performed for pairs of genetic variants showing signatures of antagonistic pleiotropy for disruptive adaptation to ecological selection for top or bottom in allopatric populations. This included variants *II:1718756* (with the derived allele A in high frequency in bottom populations relative to the reference allele present in high frequency in top populations) and *II:1741521* (derived allele T in high frequency in top populations relative to reference allele in high frequency in bottom populations). These two loci were identified in populations with the α genetic background, located 22,765 bp apart. For populations derived from the β genetic background, we performed the same analysis on variants *I:2319886* (with the derived allele T in high frequency in bottom populations relative to the reference allele present in high frequency in top populations) and *I:2319922* (derived allele T in high frequency in top populations relative to reference allele in high frequency in bottom populations). This second pair of loci was 36 bp a part.

Given the close physical proximity between these pairs of variants, the strong correlation in allele frequency of variants could be due to ecologically mediated selection acting 1) on both variants either in the opposite or same direction generating linkage disequilibrium (hypothesis 1), or 2) on only one of the two variants dragging along a physically linked neutral locus (hypothesis 2).

Sequencing of population pools did not allow haplotype inference. Yet, the initial allele frequencies of both loci in the ancestral populations restrict the range of possible initial haplotype frequencies. In the ancestral α population, we observed an allele frequency of 0.5 and 0.2 for the derived alleles *II:1718756_A* and *II:1741521_T*, respectively (or 0.5 and 0.8 for the reference alleles). We ran deterministic simulations under hypothesis 2 where only one of the locus was under selection. In the first case, the variant *II:1718756* was under selection (A allele being beneficial under bottom selection) and the other locus *II:1741521* was neutral. The initial frequencies of haplotypes A1B1, A1B2, A2B1, and A2B2 were denoted as X1, X2, X3 and X4. A1 represents the allele under selection (II:1718756_A in this case) with an alternative reference allele A2; B1 and B2 were assumed to be neutral. This translated into a fitness matrix such as ωA1B1 = ωA1B2 = 1 and ωA2B1 = ωA2B2 = 1 - s, where *s* is the selection coefficient. From the fitness matrix, marginal fitness for haplotype (*i*) over other haplotypes (*j*) and mean fitness were calculated for the population as: 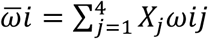 and 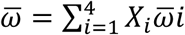. Initial haplotype frequencies were estimated considering the possible range of values such as: *X*1 + *X*2 = *f*(*A*1) = 0.5, *X*1 + *X*3 = *f*(*B*1) = 0.8, and *X*1 + *X*2 + *X*3 + *X*4 = 1. After one cycle of sexual reproduction the change in haplotype follows as:

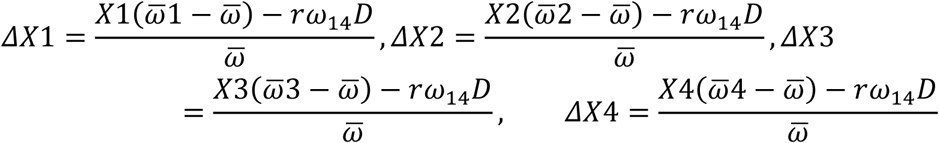

where *r* is the recombination rate (from 0 to 0.5) between the selected and the neutral loci, used also as a measurement of genetic distance, and *D* is linkage disequilibrium given by: *D* = *X*1*X*4 − *X*2*X*3. Decrease of *D* was modelled as a function of *r* over time: *D*^′^ = (1 − *r*)*D*.

This model was used to run simulations with different parameters for X1, *r* and *s*. Final allele frequencies for the neutral locus were reported once the selected locus reached fixation (X1 + X2 > 0.999). This was done for both selective regimes, one in which A1 was positively selected (allopatric bottom populations) and another one in which A2 was under positive selection (allopatric top populations). We then considered the opposite case, when the variant conferring a selective advantage was under ecological top selection (allele *II:1741521_T* positively selected in top populations) and the second locus was neutral (*II:1718756*). We compared the predicted range of allele frequencies when variants were assumed to be neutral with observed allele frequencies in the experiment. The prediction is that under physical linkage between a selective and a neutral variant (hypothesis 2), the range of observed allele frequencies should be within the simulated intervals. In the case of disruptive selection of both loci, the range of observed allele frequencies should be larger than in the simulations, since the second selected variant would increase in frequency even after the first one fixed. The same analysis was repeated using the pair of loci from the β populations *I:2319886* and *I:2319922*.

## Acknowledgments

We thank S. Lorena Ament-Velásquez, Roger Butlin, Ulrich Knief, Dirk Metzler, Claire Peart, Ricardo Pereira, Rike Stelkens, Matthias Weissensteiner, and members of the Immler and Wolf labs for providing intellectual input on the various analyses, and comments on the manuscript. We are further indebted to Tauseef Ahmad, Gaby Kumpfmüller, Hildegard Lainer and Natalia Zajac for help with laboratory work, Douglas Scofield for bioinformatics support and Ben Haller for support running *SLiM*. We further acknowledge support for genomic data generation from the SNP&SEQ Technology Platform in the National Genomics Infrastructure, Uppsala, Sweden. Flow cytometry was performed at the Core Facility FlowCyt at LMU. The computational infrastructure was provided by the UPPMAX Next-Generation Sequencing Cluster and Storage (UPPNEX) project funded by the Knut and Alice Wallenberg Foundation and the Swedish National Infrastructure for Computing. Funding was provided to JW by LMU Munich, Science of Life Laboratories National Projects and Uppsala University.

## Contributions

ST, BN, SI and JW conceived the study; ST, BN and BW performed experiments; ST and BW performed phenotypic measurements. All analyses were performed by ST with contributions from BN in phenotypic analyses. ST and JW wrote the manuscript with input from BN and SI.

## Data and code availability

All data generated for this study are archived in the sequence read archive under bioproject ID PRJNA604890 at the National Centre of Biotechnology Information (www.ncbi.nlm.nih.gov/sra). All code used for the analyses, fitness data and a list of genetic variants (.vcf format) is available at https://github.com/EvoBioWolf/SchPom_Exp_AdaptDiv and Zenodo DOI: 10.5281/zenodo.4133489 ^88^.

