## Supplementary Material for "Experimental evolution of adaptive divergence under varying degrees of gene flow"

#### Abstract

**Adaptive divergence is the key evolutionary process generating biodiversity by means of natural selection. Yet, the conditions under which it can arise in the presence of gene flow remain contentious. To address this question, we subjected 132 sexually reproducing fission yeast populations sourced from two independent genetic backgrounds to disruptive ecological selection and manipulated the level of migration between environments. Contrary to theoretical expectations, adaptive divergence was most pronounced when migration was either absent ('allopatry') or maximal ('sympatry'), but was much reduced at intermediate rates ('parapatry', 'local mating'). This effect was apparent across central life history components (survival, asexual growth, and mating), but differed in magnitude between ancestral genetic backgrounds. The evolution of some fitness components was constrained by pervasive negative correlations (trade-off between asexual growth and mating), while others changed direction under the influence of migration (e.g. survival and mating). In allopatry, adaptive divergence was mainly conferred by standing genetic variation and resulted in ecological specialization. In sympatry, divergence was mainly mediated by novel mutations enriched in a subset of genes and was characterized by the repeated emergence of two strategies: an ecological generalist and an asexual growth specialist. Multiple loci showed consistent evidence for antagonistic pleiotropy across migration treatments and provide a conceptual link between adaptation and divergence. This evolve-and-resequence experiment demonstrates that rapid ecological differentiation can arise even under high rates of gene flow. It further highlights that adaptive trajectories are governed by complex interactions of gene flow, ancestral variation and genetic correlations.**

**Supplementary Figures……………………………………..pages 2-31**

**Supplementary Tables……………………………………..pages 32-34**

### Supplementary Figures

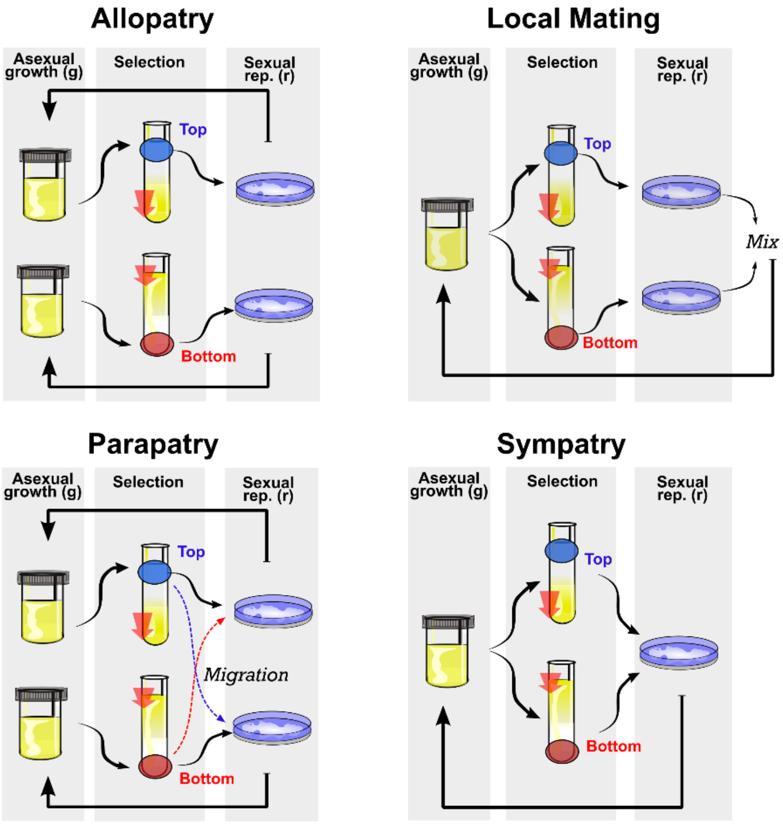


**Supplementary Figure 1.**

**Schematic illustration of the experiment.** A 6-day experimental cycle consists of asexual population growth, ecological selection and sexual reproduction. The experiment uses disruptive selection on settling speed as ecological contrast collecting cells from the Bottom (B) or the Top (T) of a column after a predefined period of time. The red arrows indicate the strength of administered gravity during ecological selection. Populations are grouped by the degree of migration experienced between top and bottom selected populations. Allopatry (Allo): 44 independent replicate populations are propagated asexually. Subsequently, half of the populations are subjected to bottom selection, half to top selection. Sexual reproduction occurs within each population. Parapatry (Para): 44 independent replicate populations are divided into 22 non-independent population pairs experiencing opposite ecological selection (top or bottom selection). After selection, 5% of the selected cells are reciprocally transferred between populations of each pair. Sexual reproduction occurs after migration within each population. Local Mating (LM): 22 independent populations are grown asexually and experience disruptive selection for both top and bottom selection. Sexual reproduction occurs in each resulting fraction independently. The spores produced from both fractions are then mixed and transferred for asexual growth. Sympatry (Sym): 22 independent populations are grown asexually and experience disruptive selection for both top and bottom selection. Prior to sexual reproduction the two fractions are fully mixed.

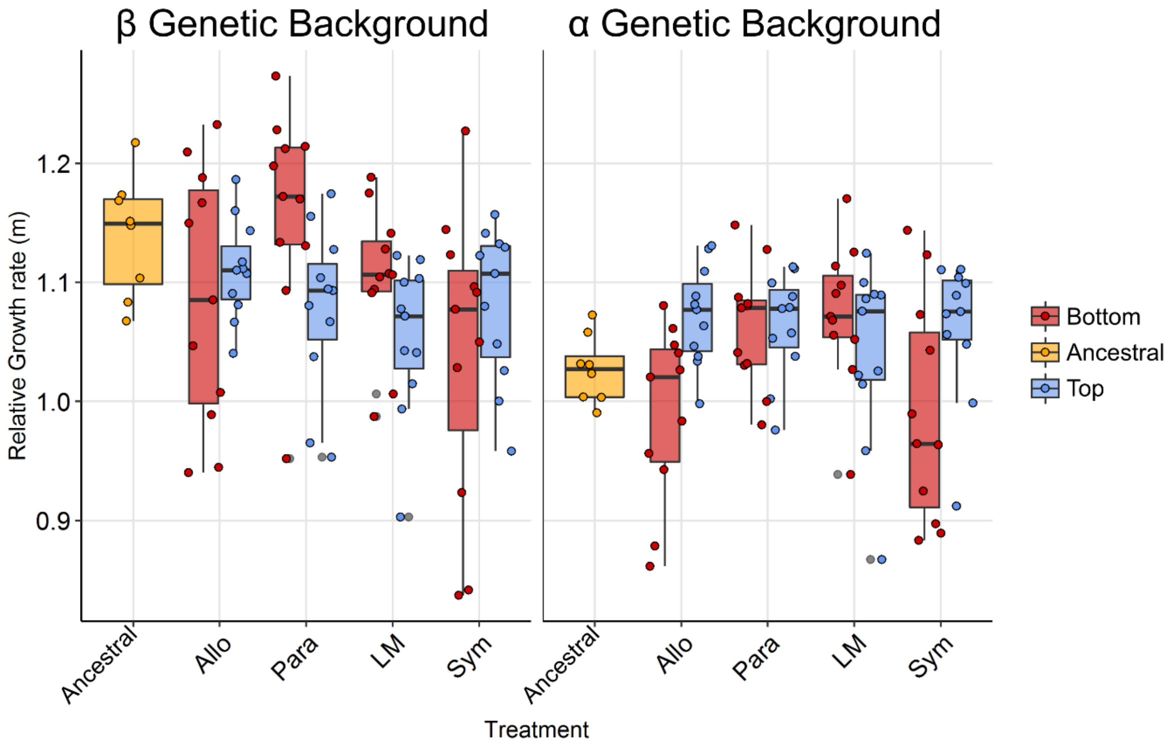


**Supplementary Figure 2.**

**Growth rate relative to fluorescent reference strain per population.** Boxplot of distribution of growth rate relative to fluorescent reference. Each point represents the median of 8 technical replicates per population. In the ancestral population, however, each point reflects a technical replicate. Note the difference in general growth performance between ancestral genetic backgrounds. The trade-off between growth and sexual reproduction, however, remains strong for both backgrounds (see **Figure 2b**, main text).

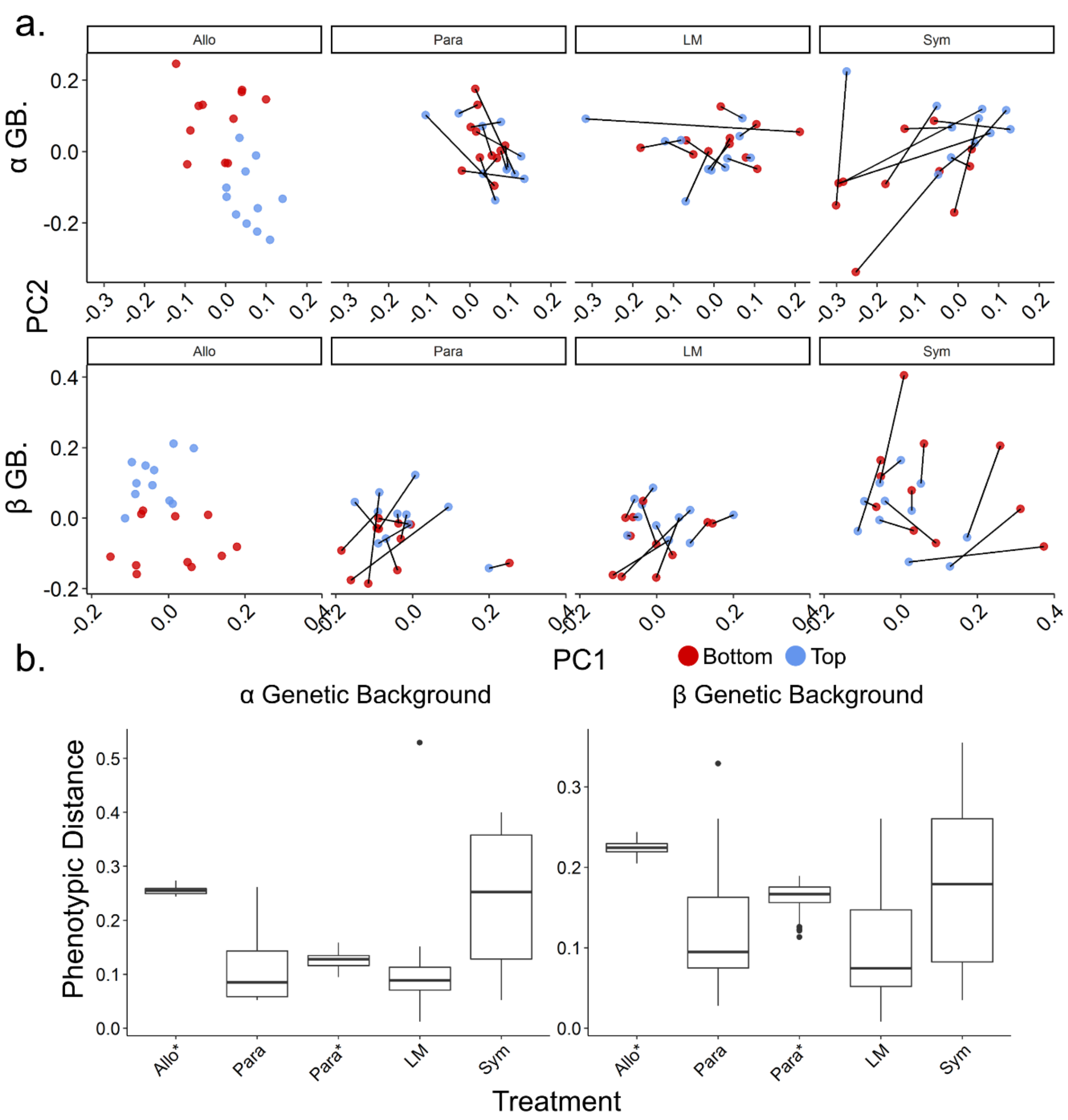


**Supplementary Figure 3.**

**Distance between relative fitness as a function of migration treatment. A.** PCA using z-score normalized fitness values per genetic background. Distribution of populations are given per treatment, with pairs of populations connected by a line. **B.** Boxplot of Euclidian distance between pairs of connected population pairs per treatment and genetic background. For allopatric populations (Allo*), the mean distance was calculated for a subset of 100 bootstrapped combinations of all top and bottom populations. In parapatric populations, the boxplot displays the distances between connected populations (Para) and bootstrapped combinations between independent top and bottom populations as in Allo* (Para*). Pairs of parapatric and local mating population are more similar in fitness than allopatric, randomized parapatric or sympatric populations. This is consistent with a homogenizing effect of gene flow for these treatments.

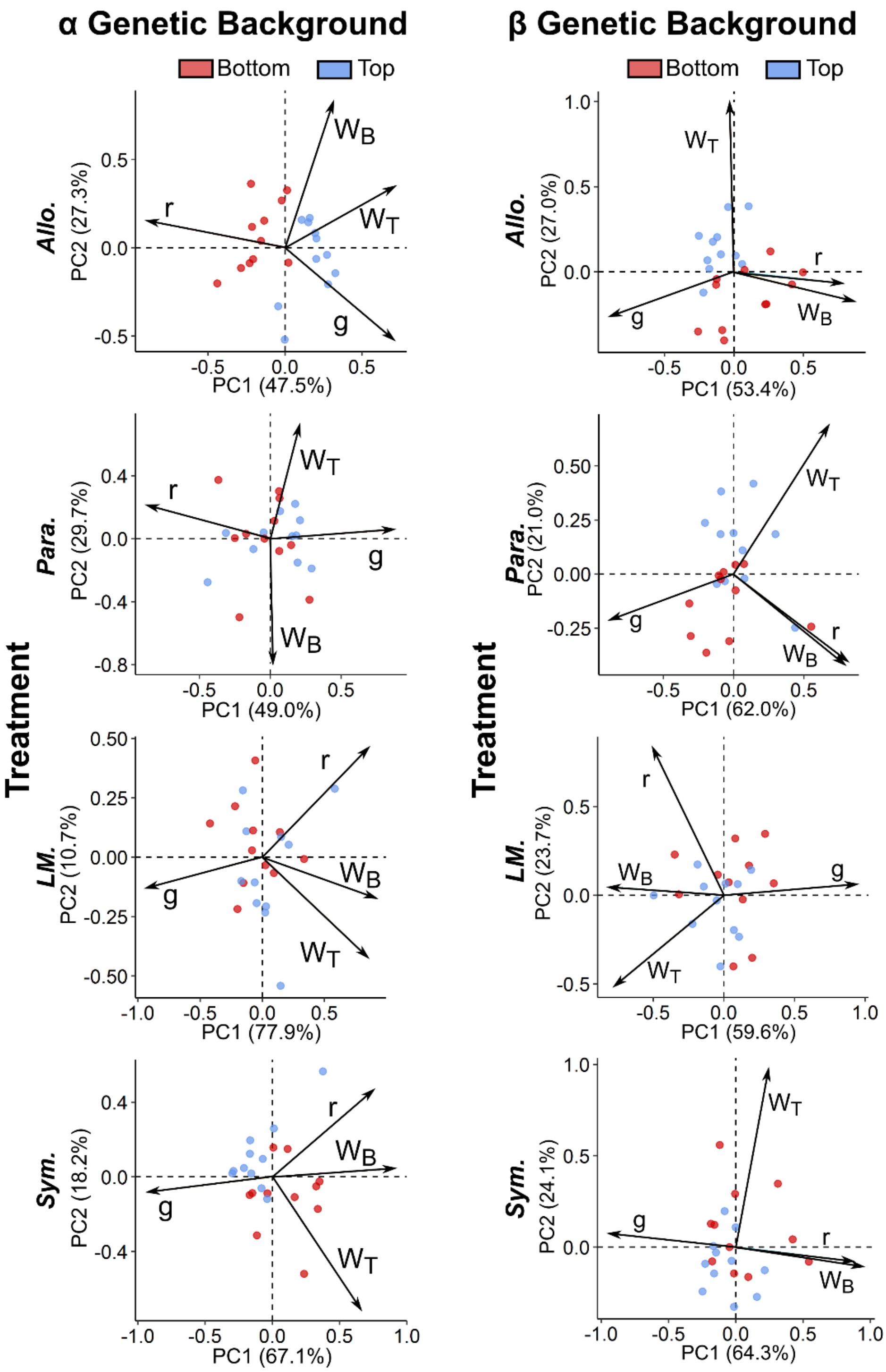


**Supplementary Figure 4.**

**Principle component analyses of z-score standardized log transformed relative fitness values across fitness components**. Each point represents one population or subpopulation subjected to the top (blue) or bottom (red) selection regime. Eigenvectors are indicated by arrows for each fitness component. Arrows pointing in the same direction reflect a positive relationship between components, arrows in opposite direction indicate a negative correlation or trade-off and orthogonal arrows suggest no relationship. The relative contribution to variance in PC1 and PC2 is symbolized by length of the arrow. Proportion of variance explained by each PC is shown in parenthesis for both axes. Results are shown for the α and β genetic background. Abbreviations of migration treatments as in the main manuscript.

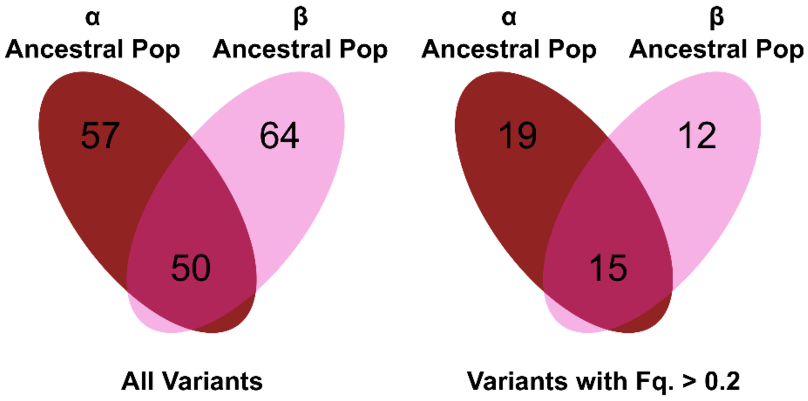


**Supplementary Figure 5.**

**Standing genetic variation**. Venn diagram displaying the number of genetic variants per genetic background in the two ancestral populations at the beginning of the experiment. The left panel includes all variants, whereas the right panel is restricted to variants with allele frequency higher than 0.2.

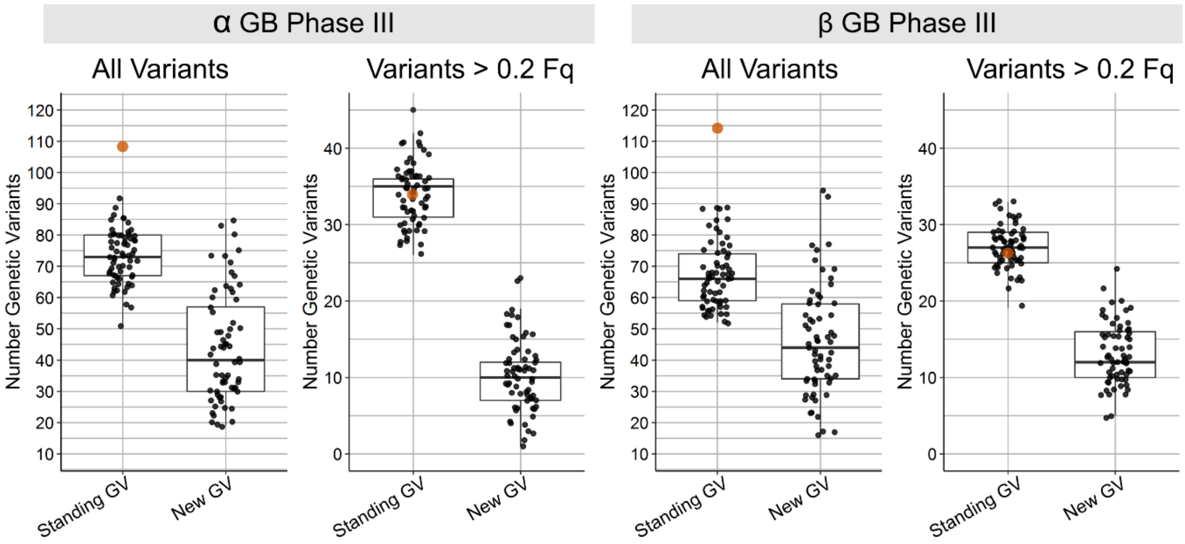


**Supplementary Figure 6.**

**Genetic variation per population**. Boxplot with number of genetic variants per population, differentiating between standing genetic variation and new mutations emerging during the experiment. Panels include either all variants or only variants with allele frequency higher than 0.2, as well as between genetic backgrounds (α and β genetic background). The orange point shows the number of genetic variants in ancestral populations at the beginning of the experiment. At the end of the experiment we counted in total 1,472 and 1,318 genetic variants (α and β respectively) of which 1,217 and 1,061 arose de-novo during the experiment. Most variants were present at low frequencies (1,179 and 1,073 variants with a maximum frequency of 0.2), and most were limited to single populations (950 and 809 variants). At the end of the experiment, each of the 132 evolved populations had between 71 and 183 mutations.

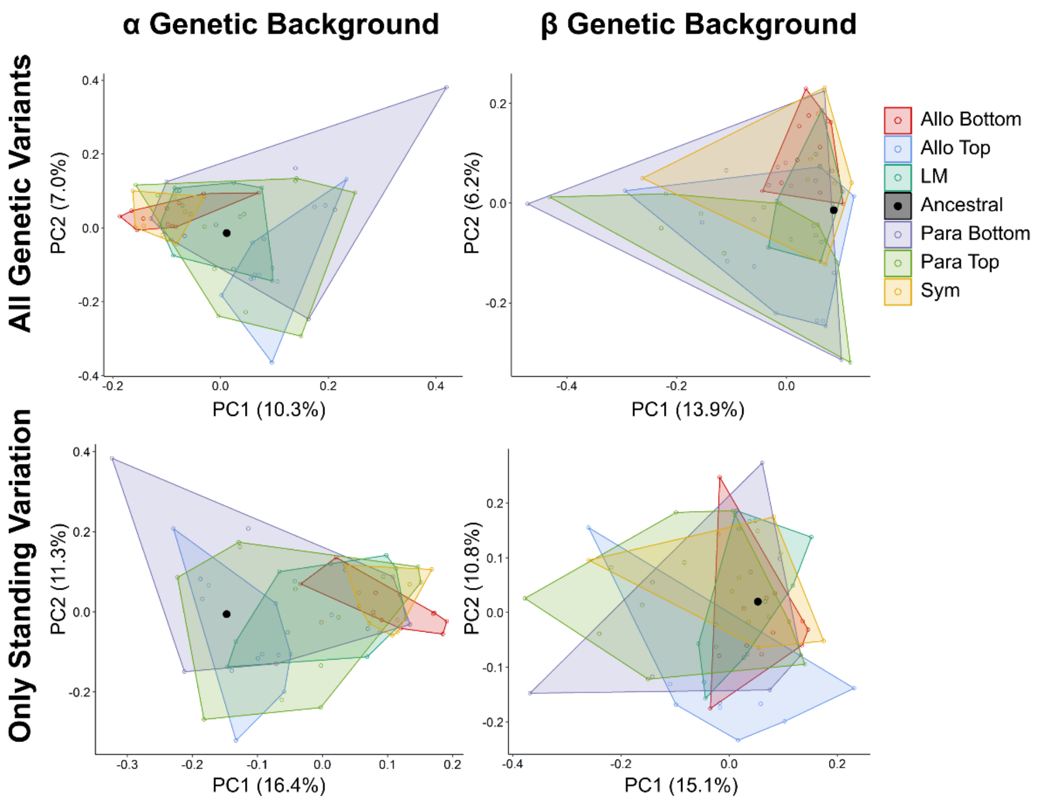


**Supplementary Figure 7.**

**Genetic differentiation between populations**. PCA including all populations per genetic background. The analysis was conducted including all genetic variants (upper graphs) or using only variants present at the beginning of the experiment (standing genetic variation; lower graphs). Decomposition of genetic variation across all populations using principal component analyses shows a similar distribution of populations for the genetic backgrounds across the different levels of gene flow for the first two principal components. Using all genetic variants, the two major axes of variation explained in total 17.3 % and 20.1 % of the genetic variation for the α and β background, respectively. This increased to 27.7 and 25.9% when using only standing genetic variation, however, the distribution of populations remained similar.

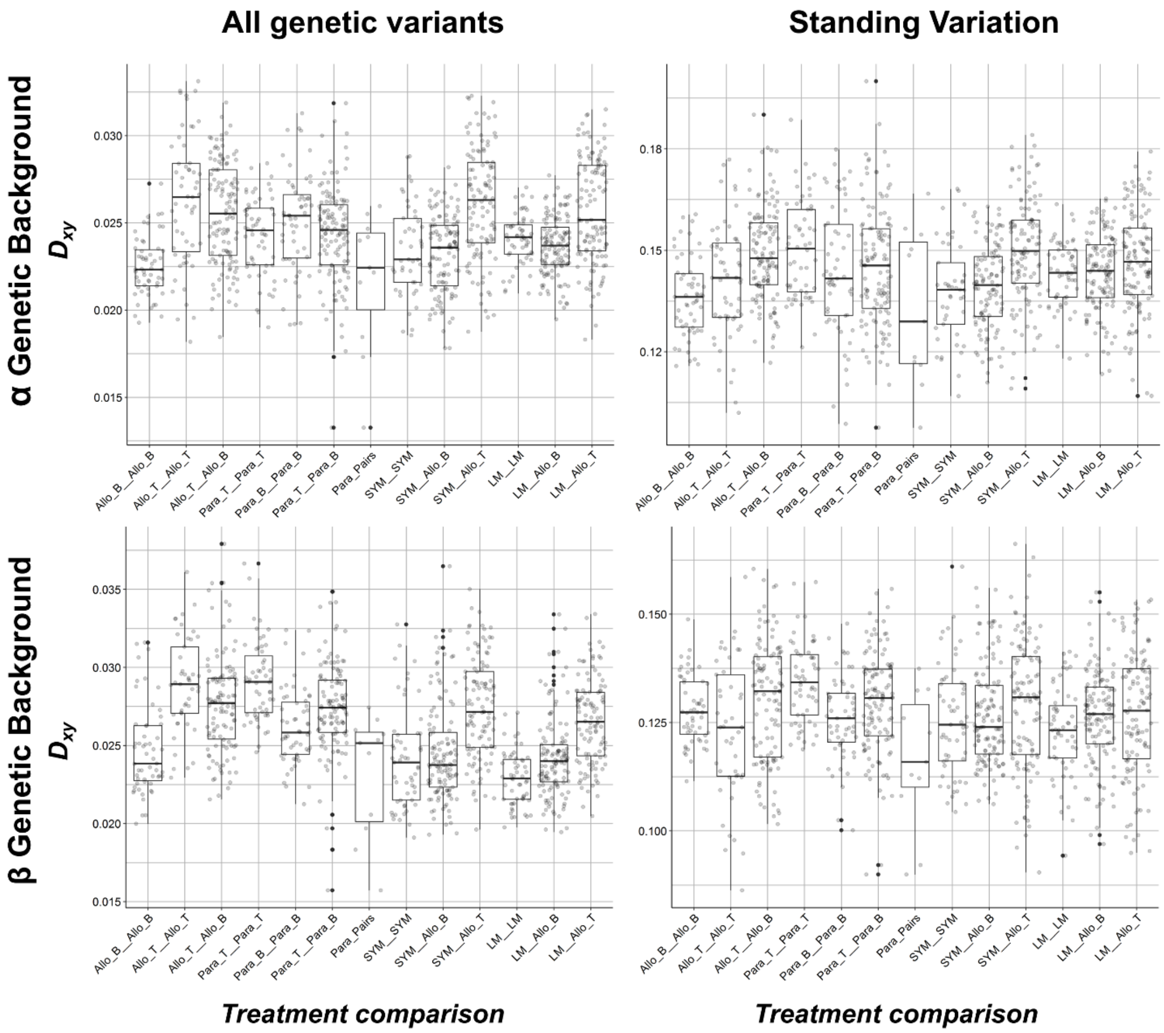


**Supplementary Figure 8.**

**$D_{xy}$ between populations.** Boxplot of distribution of $D_{xy}$ between pairs of populations. Each point represents a $D_{xy}$ value for a single pair of populations. $D_{xy}$ values were calculated using all genetic variants (left) or only standing genetic variation (right). Distribution of values were divided per genetic background (α and β). For each treatment, comparisons were done between populations with the same or opposite ecological selection regime (Top - T and Bottom – B). For parapatric populations, $D_{xy}$ was additionally calculated between connected population pairs (Para_Pairs).

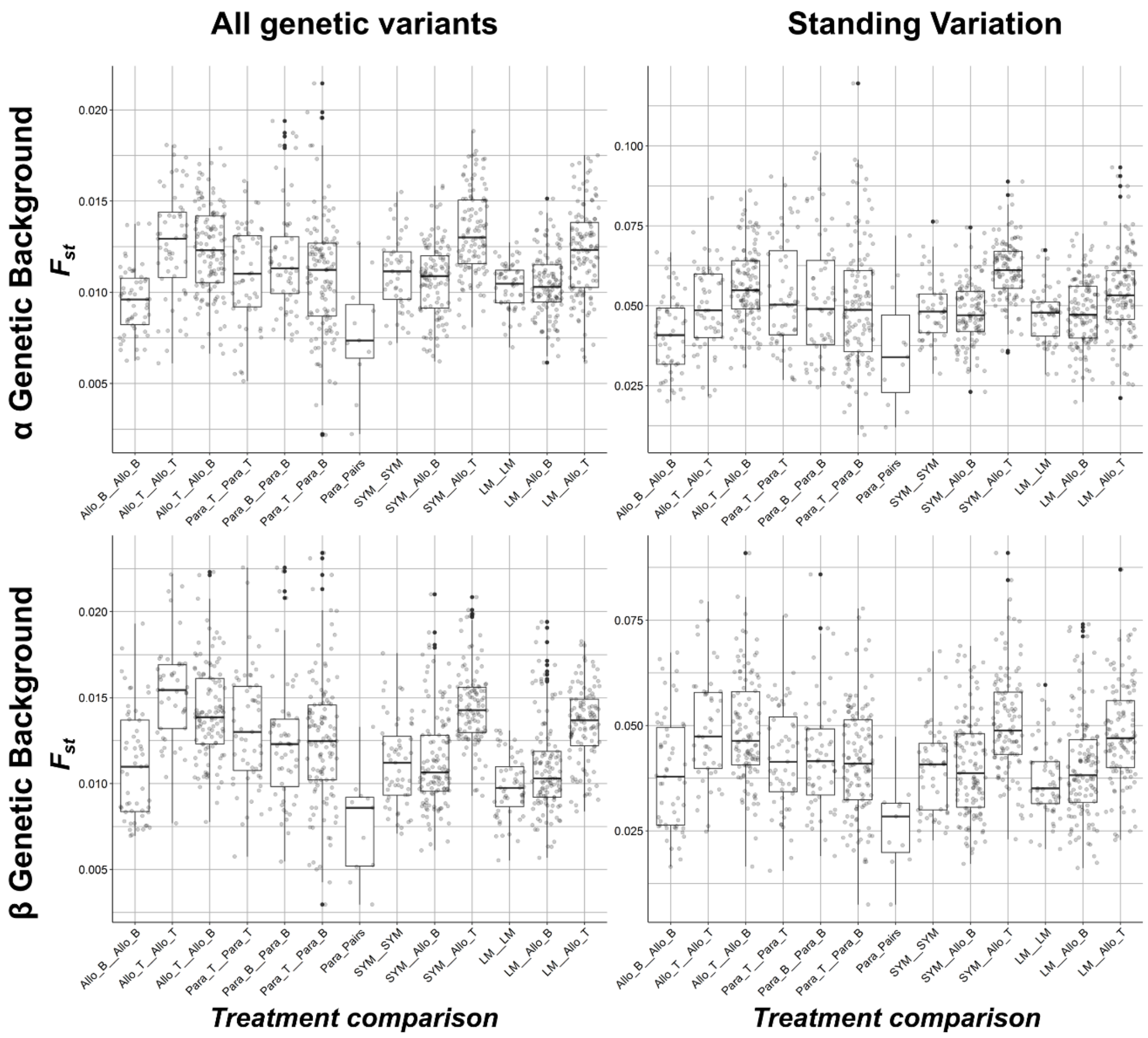


**Supplementary Figure 9.**

**$F_{st}$ between populations.** Boxplot of distribution of $F_{st}$ between pairs of populations. Each point represents an $F_{st}$ value for a single pair of populations. $F_{st}$ values were calculated using all genetic variants (left) or only standing genetic variation (right). Distribution of values were divided per genetic background (α and β). For each treatment, comparisons were done between populations with the same or opposite ecological selection regime (Top - T and Bottom – B). For parapatric populations, $F_{st}$ was additionally calculated between connected population pairs (Para_Pairs).

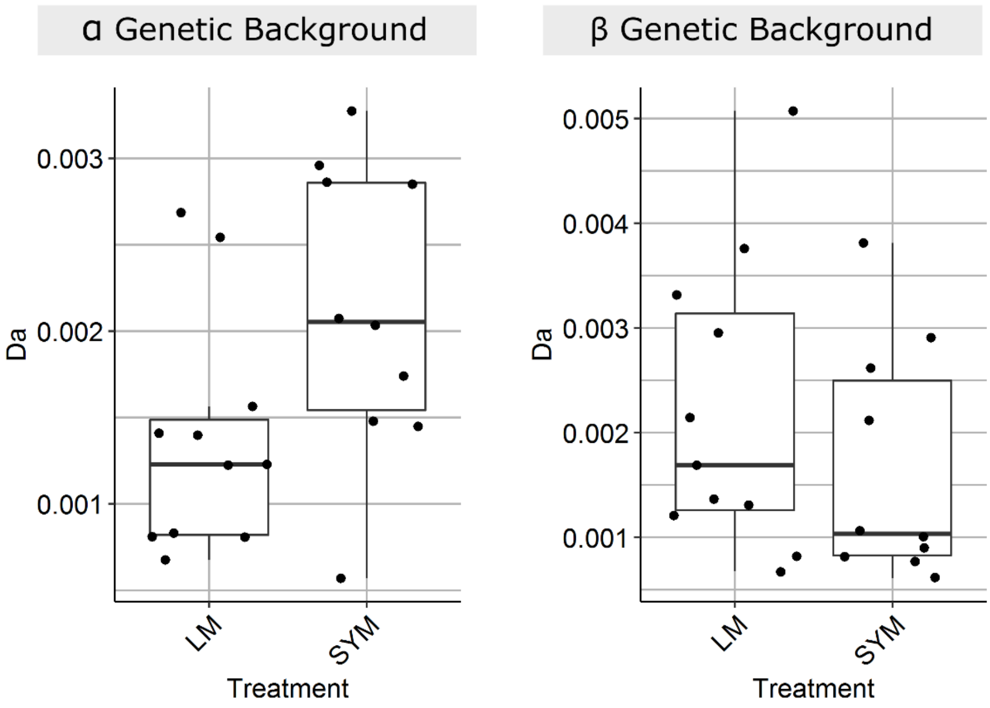


**Supplementary Figure 10.**

**Genetic divergence between Top and Bottom ecotypes.** Boxplot of genetic divergence, measured as Da (Difference between $D_{xy}$ and mean $\pi$), between the top and bottom ecotypes per population. Each point represents the comparison between top and bottom ecotypes within each population. The plot is divided by treatment (Local mating and Sympatry) and by genetic background. In the $\alpha$ genetic background genetic divergence was higher for Sympatric populations compared with Local Mating populations, which is consistent with phenotypic data. This divergence was not observed for the $\beta$ background, however the absolute phenotypic variation was lower in this genetic background (**Figure 2a**).

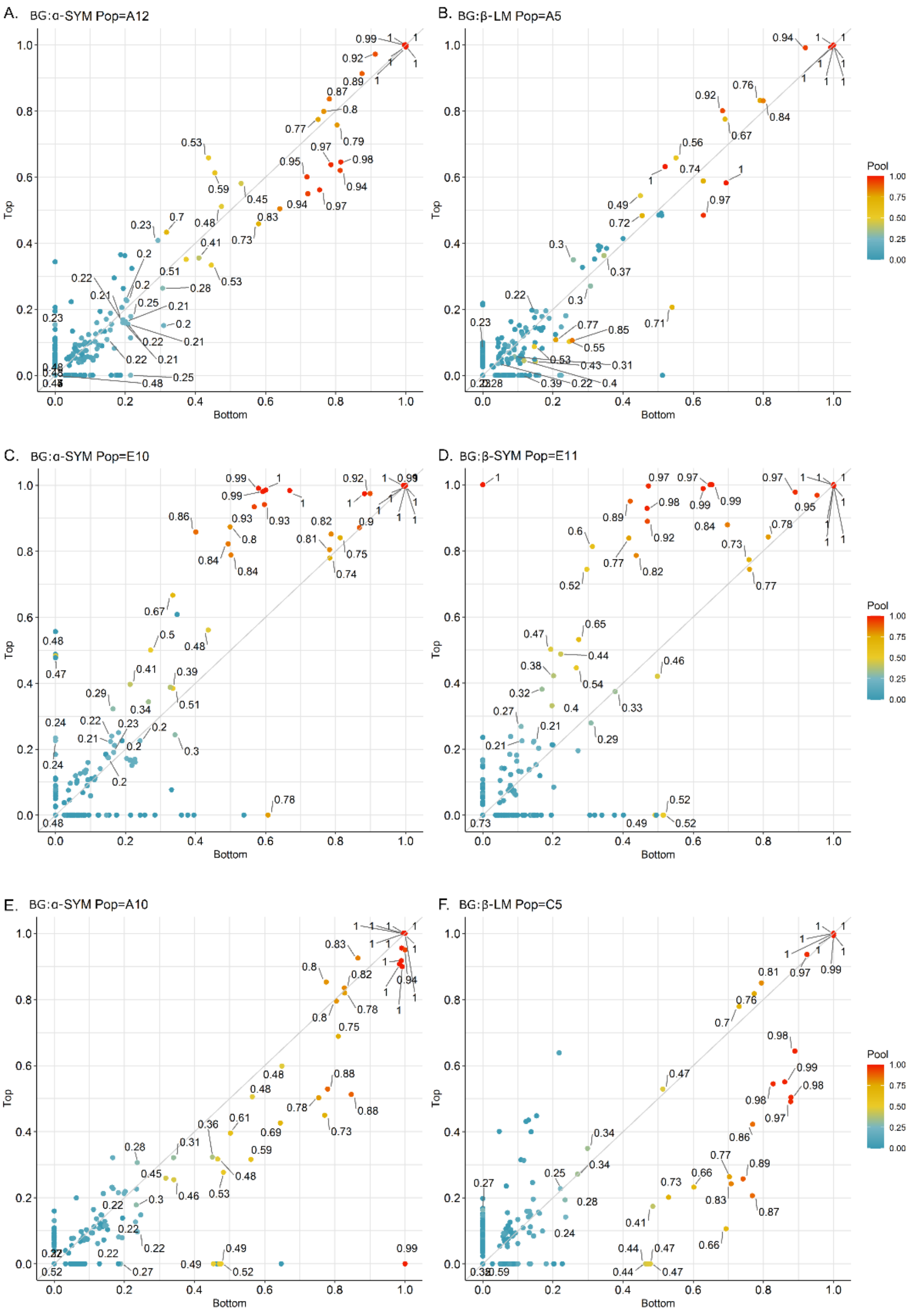


**Supplementary Figure 11.**

**Comparison between allele frequencies of the population pool, the top and bottom ecotype fractions. Examples are shown for 6 populations to represent the three possible scenarios.** Each plot shows the allele frequency for the bottom ecotype fraction (x-axis), the top ecotype fraction (y-axis) and in the original whole pool (colour gradient and labels when frequency is higher than 0.2). Each point represents one genetic variant. The different plots shows examples of populations with small differences between top and bottom fractions indicating no divergence (**A** and **B**); when the pool is more similar to the top fraction and there are larger changes in allele frequency for the bottom fraction (**C** and **D**); and when the pool is more similar to the bottom fraction and there are larger changes in allele frequency for the top fraction (**E** and **F**). This data forms the basis for **Supplementary Figure 12**.

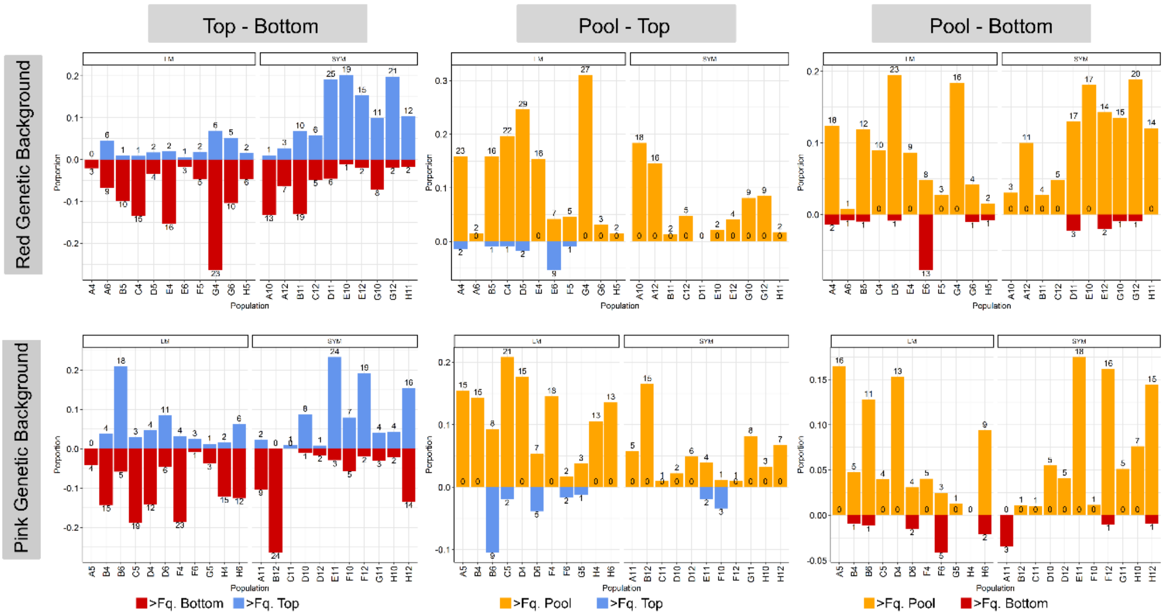


**Supplementary Figure 12.**

**Population specific statistics of allele frequency change between subpopulations.** Data are shown for the local mating (LM) and sympatry (SYM) treatment and are displayed for both ancestral genetic backgrounds (α and β). For each of the 11 populations from a treatment the allele frequencies of segregating variants were compared between subpopulations including the following comparisons. *Left panel (Top-Bottom):* Comparison between the top and bottom ecotypic fractions; comparison between the population pool (prior to ecological selection) with the top (*Middle panel: Pool-Top)* or bottom ecotypic fraction (*Right panel: Pool -Bottom).* For each population the proportion (bars) and total number (numbers above bars) of genetic variants with an allele frequency shift of > 0.2 (between the two fractions, or between the fraction and the pool) are shown. Differences are shown as absolute changes and the direction is indicated by the colour of the bar: blue indicates higher frequency for the top fraction, red higher frequency in the bottom fraction and yellow higher frequency in the pool.

For example, for the α genetic background, we found that sympatric populations showed higher similarity between the whole pool sample and the top ecotype fraction as evidenced by lower frequency shifts after top selection (lower proportion of variants differing between top and the whole pool sample (top middle panel) than between bottom and the whole pool sample  (top right panel)). In local mating populations, the patter was opposite showing higher similarity with the bottom ecotype fraction. A summary of the comparison is shown in **Supplementary Figure 13**.

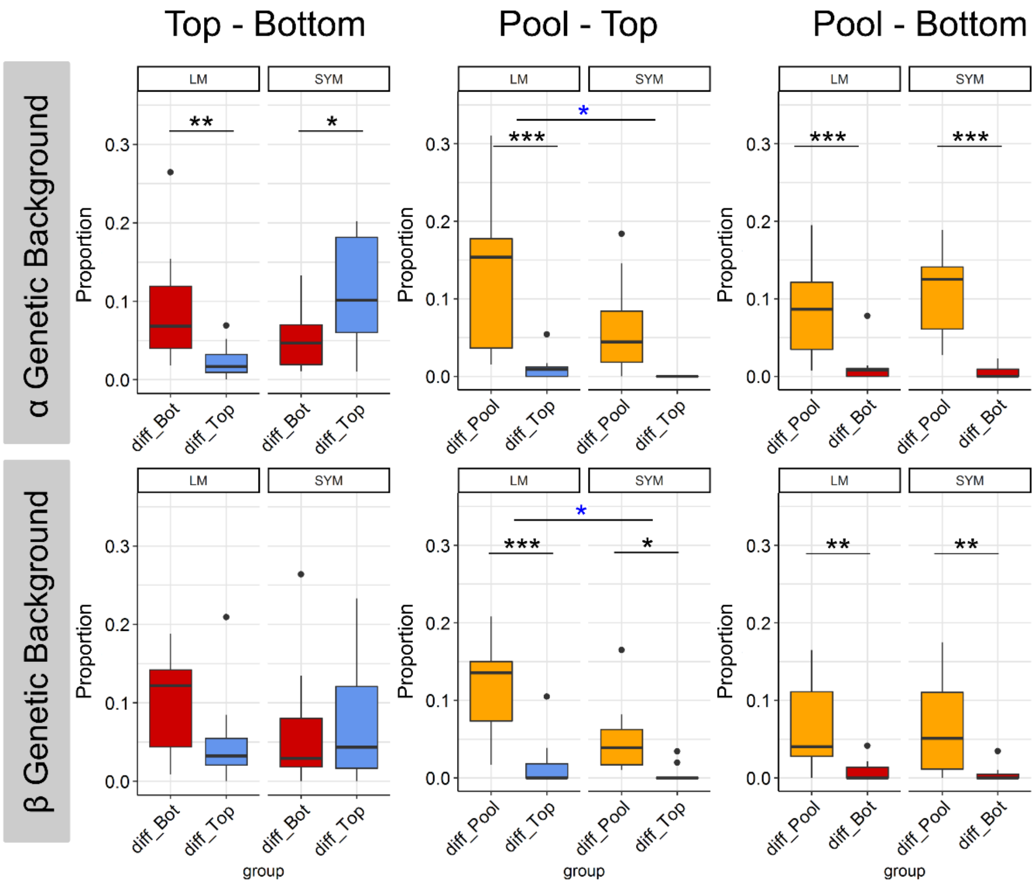


**Supplementary Figure 13.**

**Summary statistics of allele frequency change between subpopulations.** Data are shown for the local mating (LM) and sympatry (SYM) treatment and are displayed for both ancestral genetic backgrounds (α and β). For each of the 11 populations per treatment the allele frequencies of segregating variants were compared between subpopulations including the following comparisons. *Left panel (Top-Bottom):* Comparison between the top and bottom ecotypic fractions; comparison between the population pool (prior to ecological selection) with the top (*Middle panel: Pool-Top)* or bottom ecotypic fraction (*Right panel: Pool -Bottom).* For each population the proportion of genetic variants with an allele frequency shift of > 0.2 were calculated. The resulting proportion of genetic variants changing in either direction as displayed for each population in **Supplementary Figure 12** are here summarized to show the general trend. Following the colour scheme from **Supplementary Figure 12** boxplots are coloured by the fraction showing an increase in frequency, i.e. blue for the top ecotype (diff_Top), red for the bottom ecotype (diff_Bottom) or yellow for the pool (diff_Pool). In the α background we found that sympatric populations showed higher similarity between the whole pool sample and the top ecotype fraction (higher proportion of variants with increased frequency in top fraction, lower proportion of variants differing between top and the whole pool sample, and higher proportion of variants differing between bottom and the whole pool sample). In local mating populations, the pattern was opposite showing higher similarity with the bottom ecotype fraction. In the β background, although there was a similar trend, the difference between treatments was lower which is consistent with no observed divergence from the phenotypic data. Significance of the difference between groups was tested using a quasibinomial model in a nested generalised lineal model with treatment and fraction as fixed variables. Significant difference between treatments and group are shown with blue and black asterisks respectively.

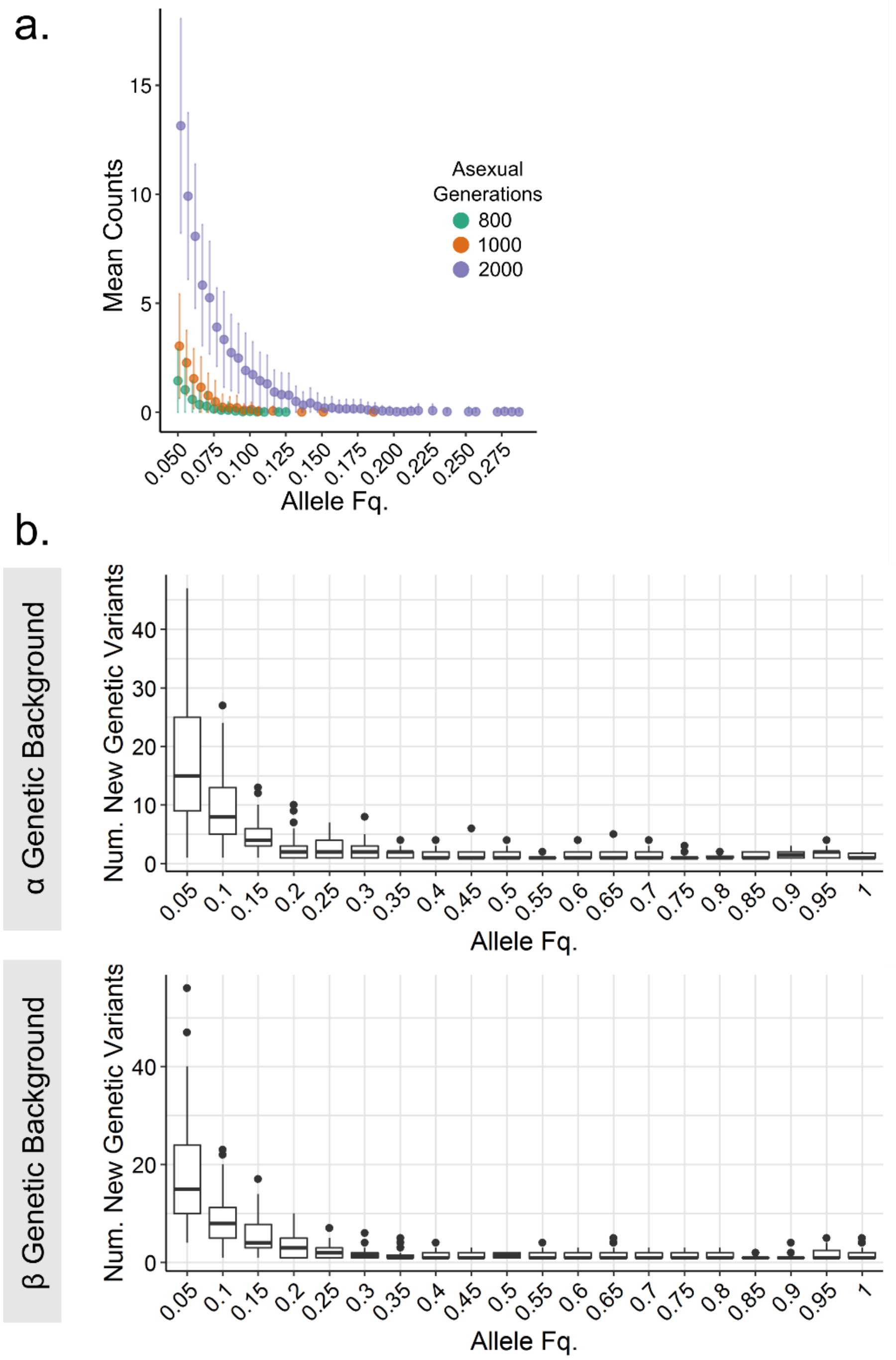


**Supplementary Figure 14.**

**Simulations of neutral variants and site frequency spectra on novel genetic variants in phase III. A.** Individual based forward simulations including only neutral variants. Distribution of number of genetic variants from 100 simulations (mean values with points) per allele frequency. Vertical lines show the standard deviation. Simulations were run in cycles until 800 to 2000 asexual generations were reached (colour). In the experiment, we estimated 53 cycles corresponding to around 700 generations. Variants with allele frequencies below 0.05 are not shown. After 800 generations of simulation variants did not exceed frequencies of 0.13, and even after 2,000 generations remained below 0.3. **B.** This is in stark contrast to 21% and 28% (in the α and β backgrounds, respectively) of novel mutations observed in the empirical data having frequencies of above 0.25. The boxplots show the empirical distribution of site frequency spectra per population after evolution, for the α and β genetic background. The figure only includes novel genetic variants emerging during the experiment without considering standing genetic variation. Note different scales on y-axis for the α and β genetic background.

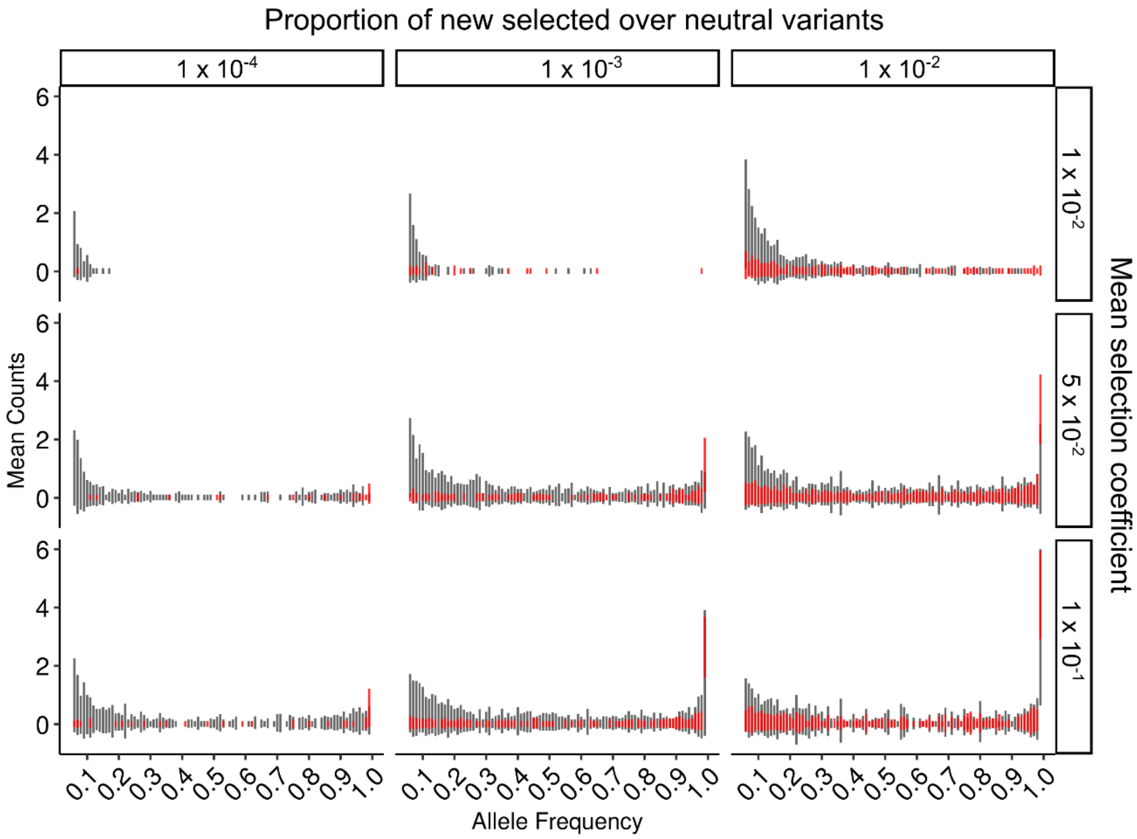


**Supplementary Figure 15.**

**Simulations of neutral variants and selected variants.** Individual based forward simulations including neutral variants (black bars) and selected variants (red bars). Distribution of the number of genetic variants from 100 simulations per allele frequency. Vertical lines show the standard deviation. Variants with allele frequencies below 0.05 are not shown. Panels show simulations using different ratio of novel, selected over neutral variants from 100 to 10000 neutral mutations per selected variant, and different mean selection coefficient from an exponential distribution. These simulations suggest that empirical frequencies of > 0.25 are readily obtained for targets of selection or linked sites assuming mean selection coefficients in the range of (s=0.001-0.1) and target densities of 1 in 100-10000 neutral SNPs.

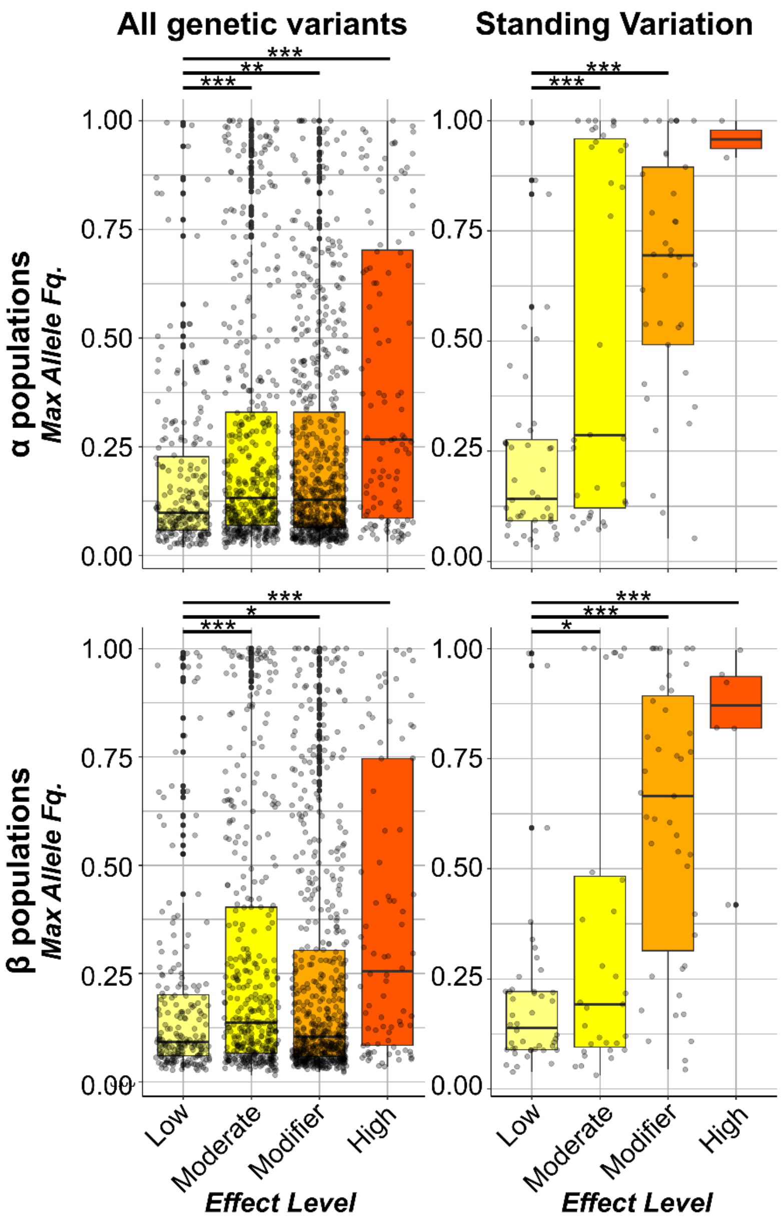


**Supplementary Figure 16.**

**Distribution of effect sizes for all genetic variants.** Boxplot of distribution of maximum allele frequencies grouping variants by the predicted functional effect level. Each point represents the maximum allele frequency observed in all populations per variant. Upper and lower panels are differentiated by genetic background (α and β populations) and include all variants (left) or only variants already present in the respective ancestral population (standing genetic variation, right). Effect levels as described in method section. P-value < 0.001 (***), < 0.01 (**), or < 0.05 (*). For further details see methods.

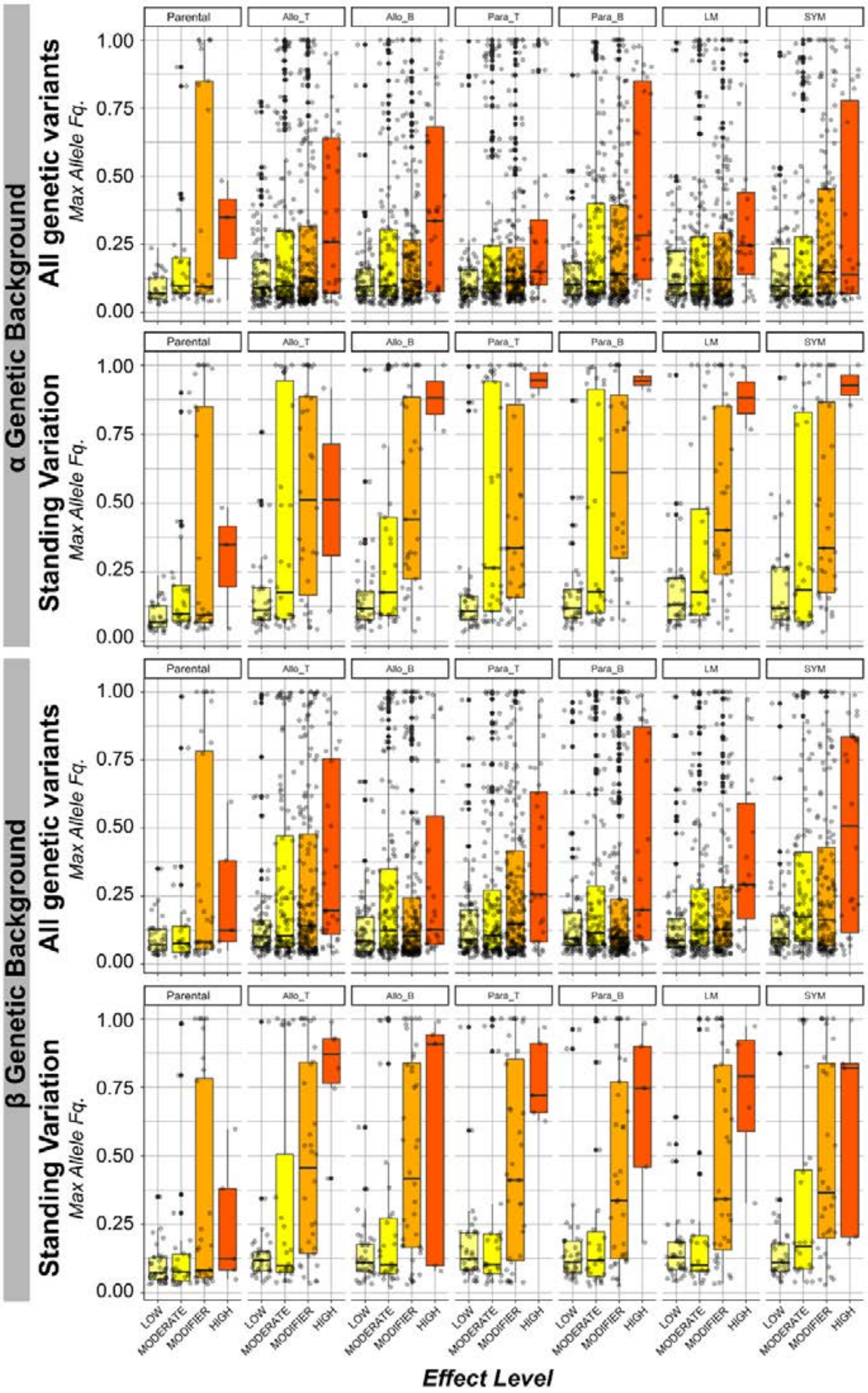


**Supplementary Figure 17.**

**Distribution of effect sizes for all genetic variants per treatment.** Boxplot of distribution of maximum allele frequencies grouping variants by the predicted functional effect level per treatment. Each point represents the maximum allele frequency observed in all populations per variant within each treatment. Evolved and ancestor populations are contrasted. Upper and lower panels are differentiated by genetic background (α and β populations) and include all variants (top) or only variants already present in the respective ancestral population (standing genetic variation, bottom). For each variant the highest effect level is used (for further details see methods). Effect levels as described in method section.

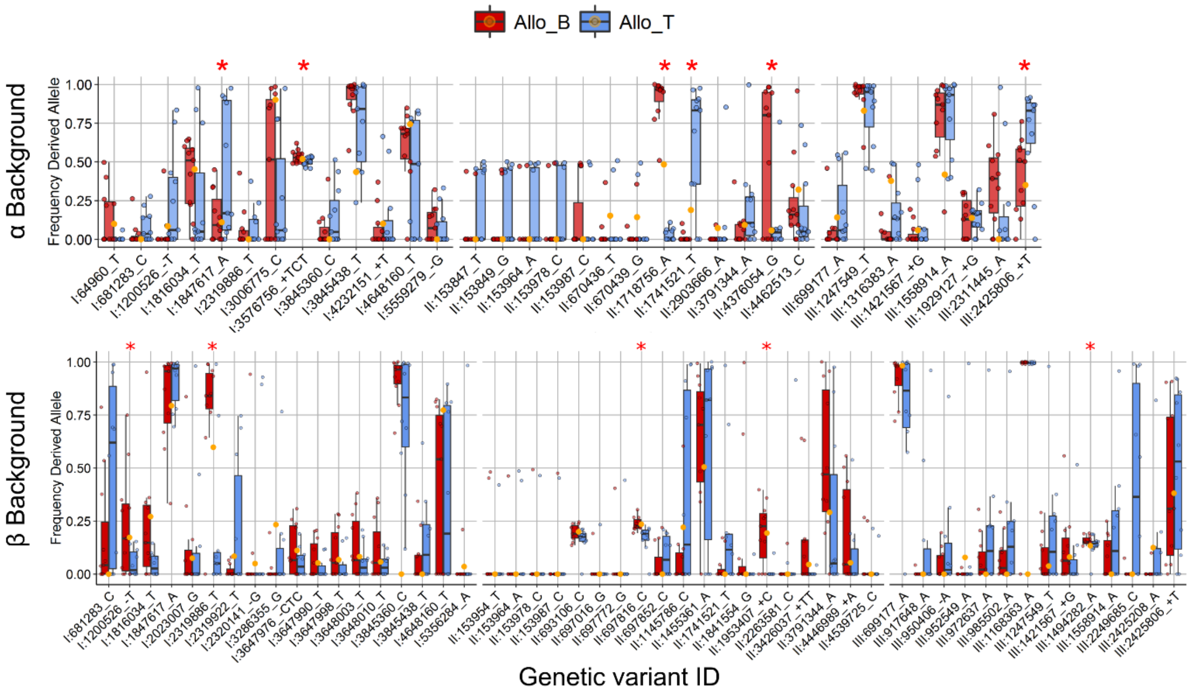


**Supplementary Figure 18.**

**Candidate genetic variants under disruptive selection per genetic background.** Boxplot of allele frequency distribution. Only variants with significant difference between top and bottom allopatric population following from logistic regression are shown. Genetic variants are labelled with chromosome, position and alternative allele relative to the reference genome. Red stars indicate variants with a significant difference between top and bottom populations, using a quasibinomial model accounting for over dispersion in the data. Allele frequencies for ancestral populations in each genetic variant is shown with yellow points.

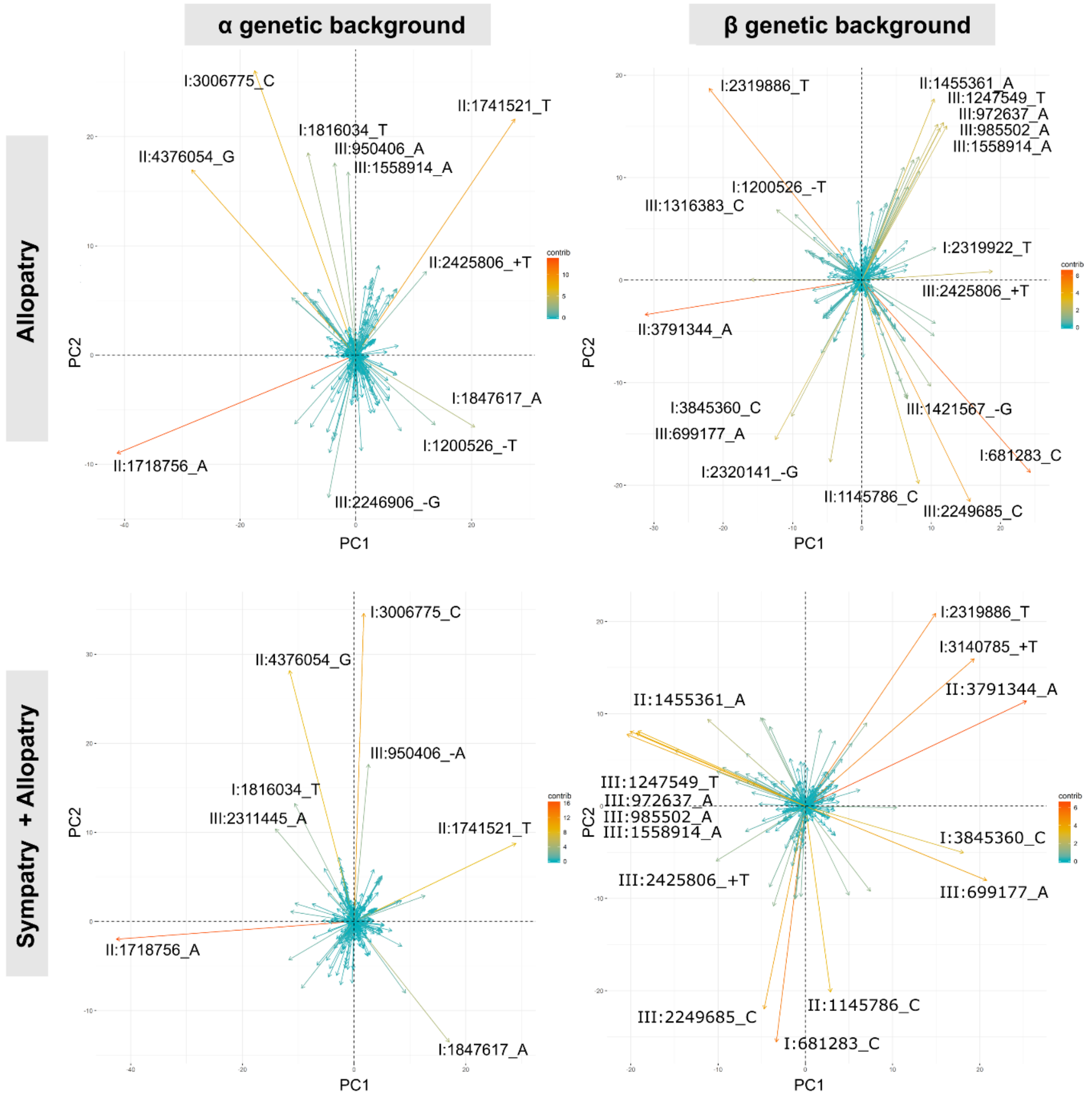


**Supplementary Figure 19.**

**Contribution to genetic variance per single nucleotide variant in PCA.** PCA plots for allopatric α and β populations. Arrows and colours represent direction and contribution per variant to total genetic variation as presented in **Figure 3a** of the main text. Genetic variants are labelled with chromosome number, position and alternative allele relative to reference. As an example of similar variants contributing to the distribution in other treatments, PCA from sympatric and allopatric populations is shown.

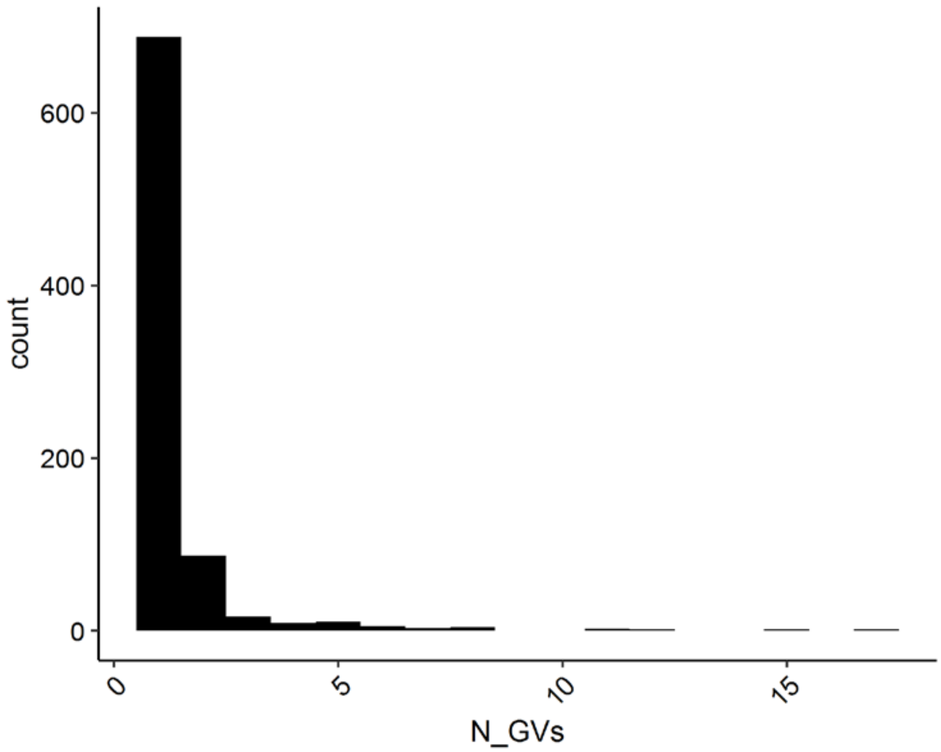


**Supplementary Figure 20.**

**Number of genetic variants per gene.** Histogram with number of genetic variants per gene combining all populations. For this plot, variants from both genetic backgrounds were pooled. To improve visualisation, in this plot the gene *SPBPJ4664.02* with 187 variants was not included.

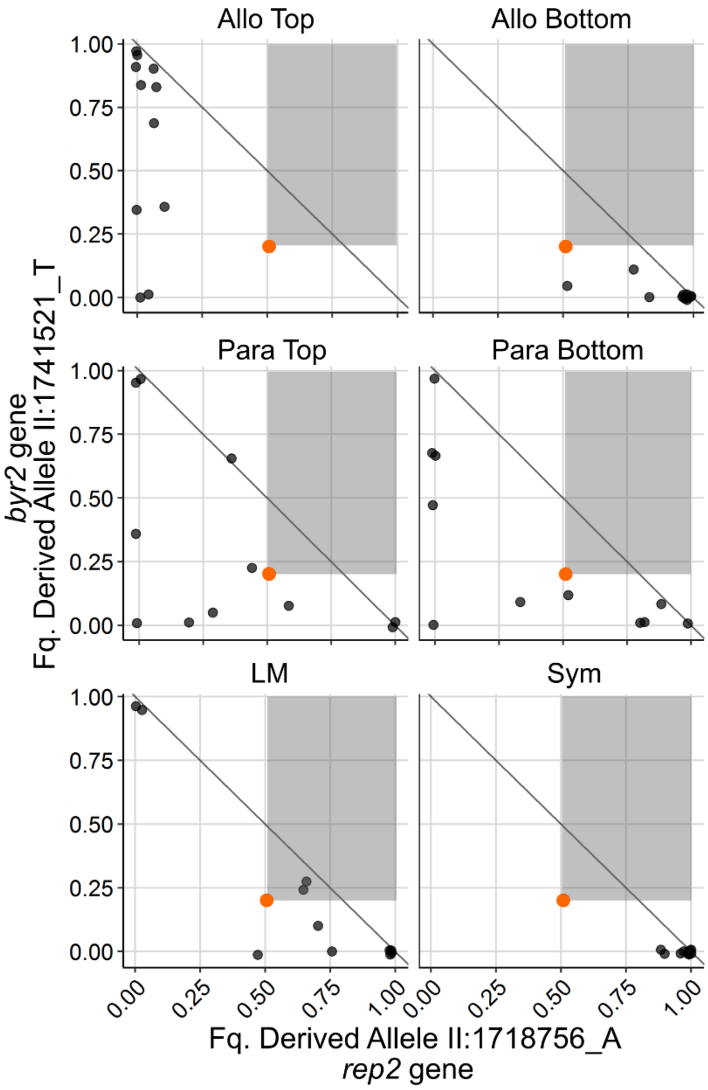


**Supplementary Figure 21.**

**Negative correlation suggesting antagonistic pleiotropy between two loci in populations of α ancestry.** Example of a pair of allelic variants from two loci that suggest antagonistic pleiotropy. Each point shows the final allele frequency for the two loci per evolved population. Ancestral allele frequencies are shown with the orange point. Panels are divided by treatment. As in **Figure 4a**, the derived allele for each locus increases consistently with ecological selective conditions in allopatric populations. Other treatments show variable end points, but populations with high frequency of derive allele at both loci are never observed. The grey area indicates allele frequency combinations that would arise under a scenario of positive selection for both derived alleles; allele frequency combinations above the diagonal line provide unequivocal evidence for coupling of both derived mutations on a single haplotype.

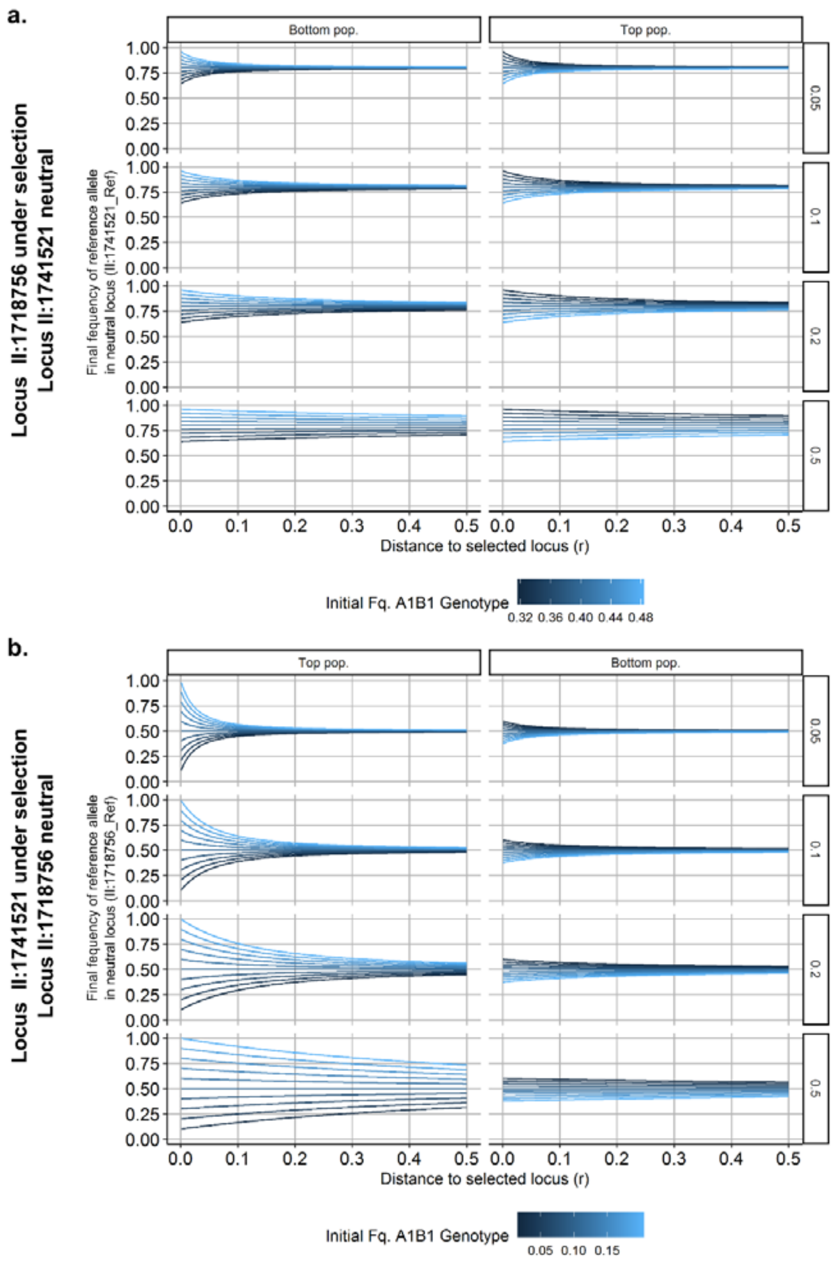


**Supplementary Figure 22.**

**Simulation of allele frequency in a pair of linked variants for populations of α ancestry.** A pair of loci from α genetic background is evaluated. Based on initial allele frequencies for both loci (frequencies in ancestral α population), the range of possible initial genotype frequencies is estimated (blue legend showing one genotype frequency). One locus is assumed to be under selection and the second one to be neutral. The figure shows the final allele frequency of the neutral reference allele once the locus under selection is fixed in the population. Different selection coefficients (from 0.05 to 0.5) and genetic distance (recombination distance r from 0 to 0.5) were tested. The same initial genotype frequencies are tested under both ecological conditions (top and bottom populations). **A.** Simulations when locus II:1718756 is under selection. As an example: with selection coefficient of 0.05, genetic distance of 0 (x axes in 0) and an initial genotype frequency for A1B1 of 0.5 (lightest blue line) (initial genotype frequency II:1718756_A/ II:1718756_Ref = 0.5), under bottom conditions (left panel), the neutral allele II:1718756_Ref goes to fixation (y axes close to 1). Using the same initial genotype frequencies, but under top conditions (right panel), only results in a reduction of the same neutral allele to around 0.65 from the original 0.8. **B.** Simulations when locus II:1741521 is under selection and II:1718756 is neutral.

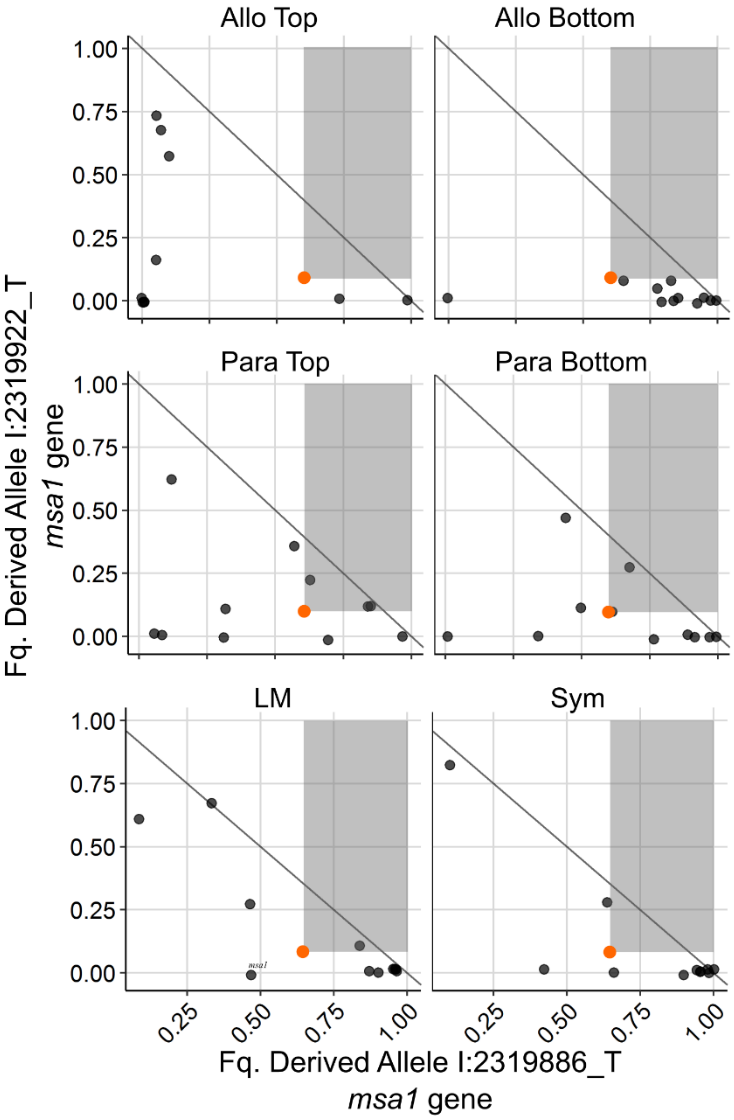


**Supplementary Figure 23.**

**Negative antagonistic pleiotropic for two loci in populations of β ancestry.** Example of a pair of allelic variants from two loci that suggest antagonistic pleiotropy. Each point shows the final allele frequency for the two loci per evolved population. Ancestral allele frequencies are shown with the orange point. Panels are divided by treatment. As in **Figure 4a**, the derived allele for each locus increases consistently with ecological selective conditions in allopatric populations. The grey area indicates allele frequency combinations that would arise under a scenario of positive selection for both derived alleles; allele frequency combinations above the diagonal line provide unequivocal evidence for coupling of both derived mutations on a single haplotype.

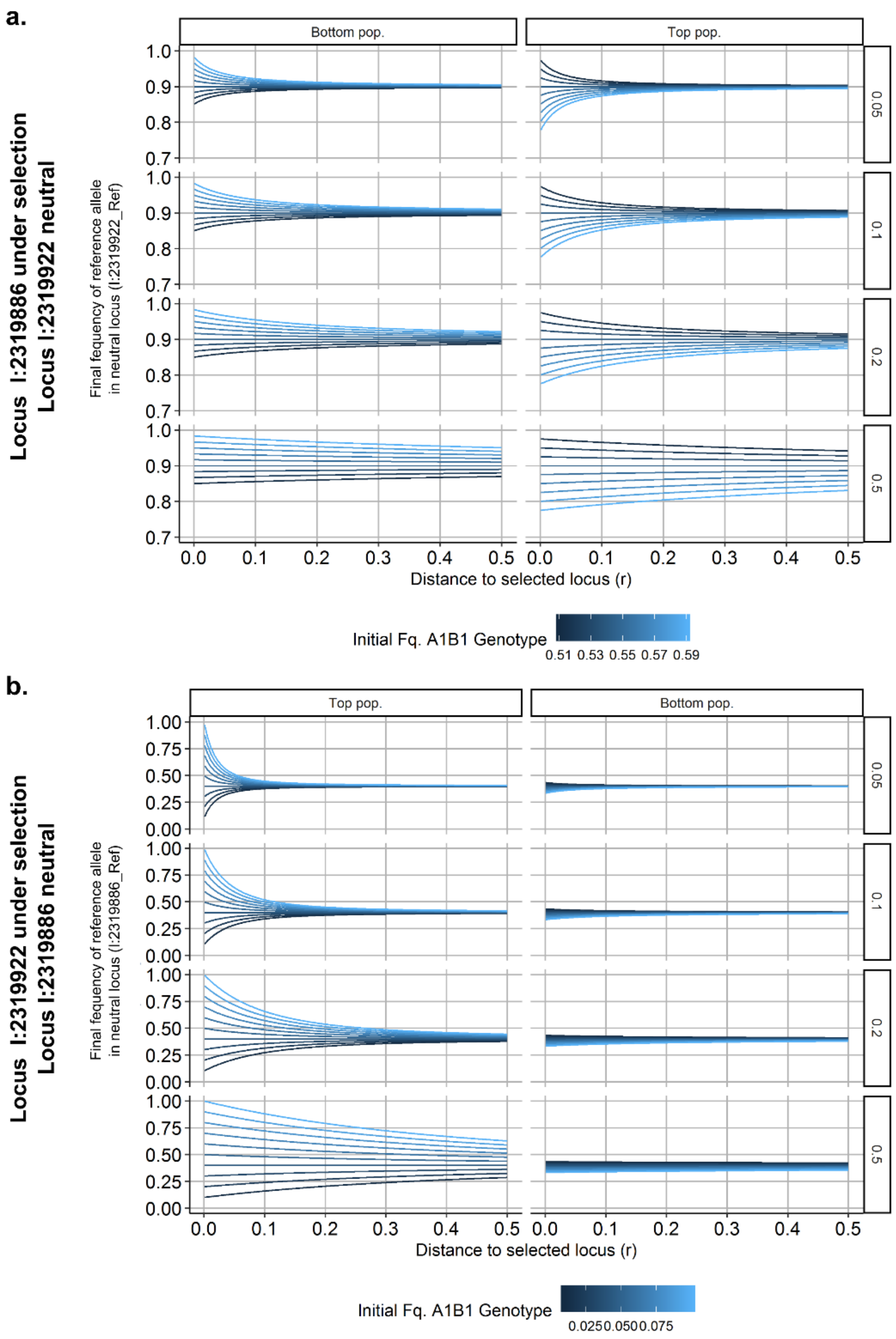


**Supplementary Figure 24.**

**Simulation of final allele frequency in pair of linked variants for populations of β ancestry.** As in **Supplementary Figure 22** but for a pair of loci from the β populations. Based on initial allele frequencies for both loci (frequencies in ancestral β population), the range of possible initial genotype frequencies is estimated (blue legend showing one genotype frequency). One locus is assumed to be under selection and the second one to be neutral. The figure shows the final allele frequency of the neutral reference allele once the locus under selection is fixed in the population. Different selection coefficients (from 0.05 to 0.5) and genetic distances (recombination distance – r from 0 to 0.5) were tested. The same initial genotype frequencies are tested under both ecological conditions (top and bottom populations). **A.** Simulations when locus I:2319886 is under selection and I:2319922 is neutral. **B.** Simulations when locus I:2319922 is under selection and I:2319886 is neutral.

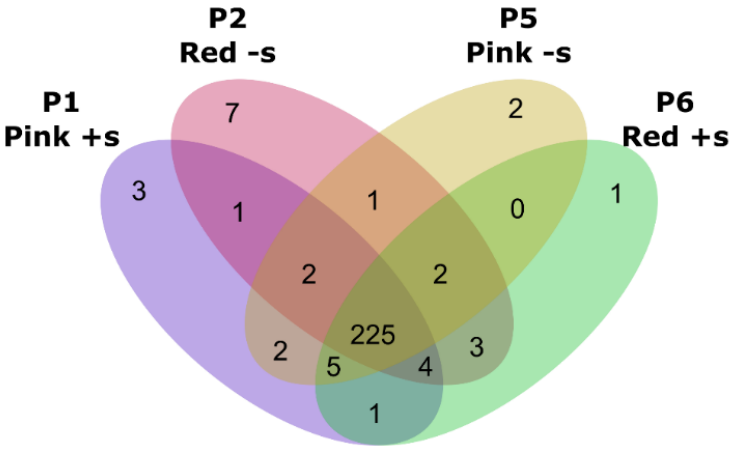


**Supplementary Figure 25.**

Number of genetic variants relative to reference *S. pombe* genome per parental strain in *phase I*. Parental strains differ in colour marker (red or pink: *ade6-M216* or *ade6-M210*) and mating type locus ($h^{+s}$ or $h^{-s}$).

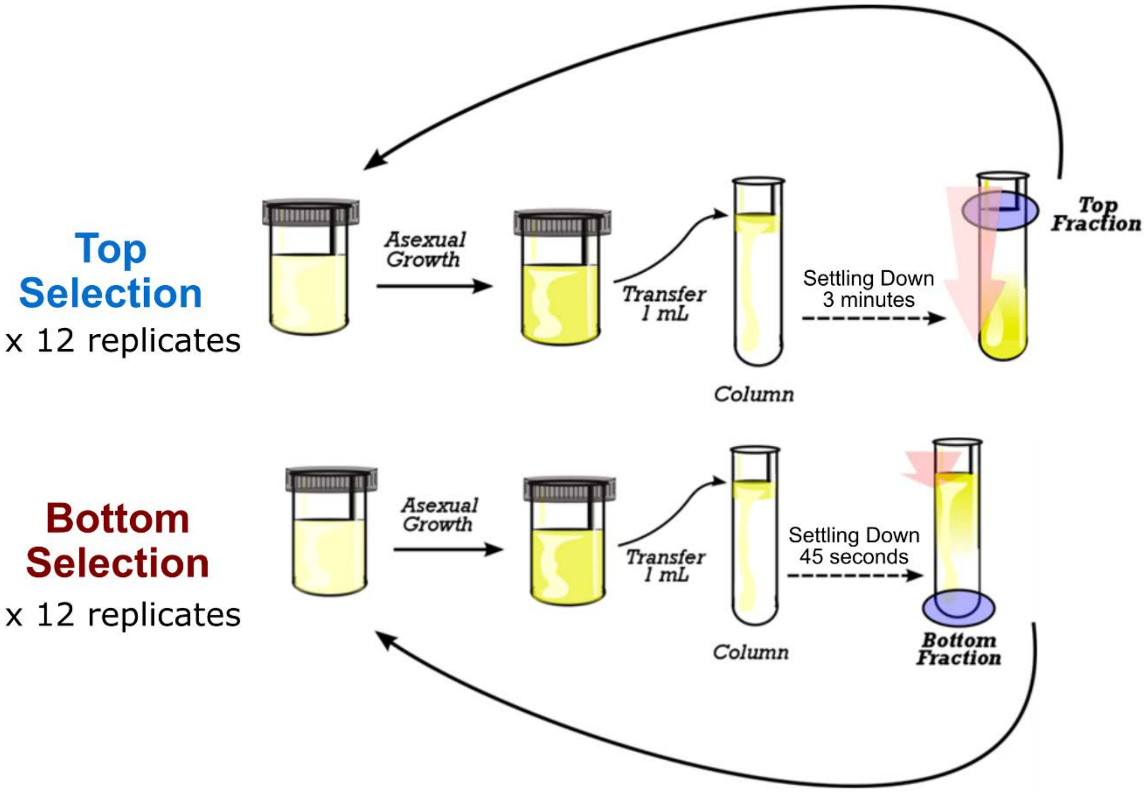


**Supplementary Figure 26.**

**Schematic overview of ecological selection during *phase I*.** The figure presents one single cycle lasting two days. During this phase, after propagating cells by asexual growth to saturation for two days, populations were selected for slow or fast settling speed (top and bottom selection, respectively). Around 1% of the total population was selected and fractions were used for a new cycle of asexual growth. The arrows illustrate the strength of gravity.

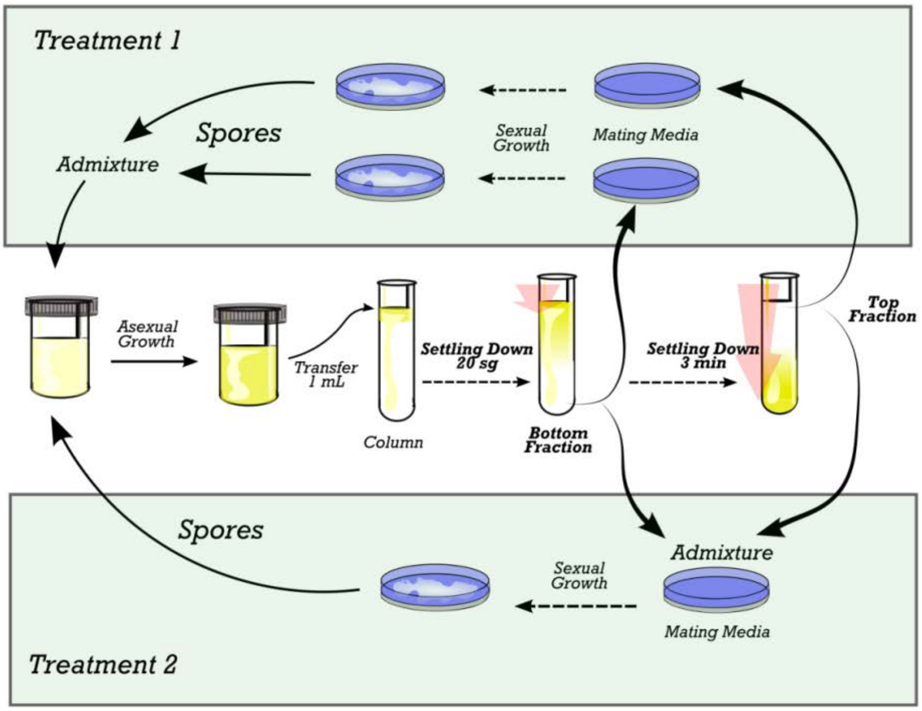


**Supplementary Figure 27.**

**Schematic overview of experimental setup during *phase II*.** The figure presents one single six days cycle, which was repeated 20 times. After asexual growth to saturation for two days, each population was selected for both slow and fast settling speed (top and bottom selection respectivelly). In treatment 1 (top pannel), each selected fraction was placed in an independent mating plate. Spores from both plates were isolated and mixed, and used to start a new cycle of asexual growth. In treatment 2 (bottom pannel), the two selected fractions were mixed before mating and placed in a single mating plate. Spores of both treatements were then mixed at equal proportion to start a new cycle. The aim of this phase was to increase mating effitiency (recombination), while maintaining ecological selection. The difference in treatment aims to ensure to maintain two top and bottom fraction (treatment 1), but also to have recombinant between selected fractions (treatment 2).

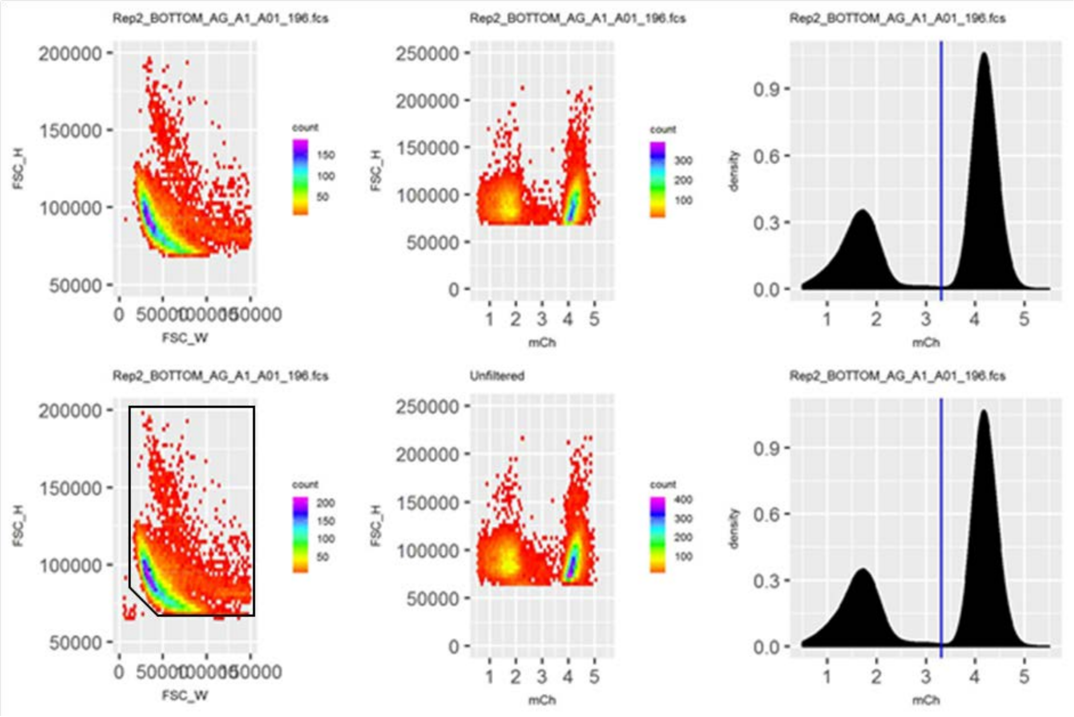


**Supplementary Figure 28.**

**Example figure for gating strategy.** Plots show before (lower panels) and after filtering (upper panels). Scatter plots show forward scatter height FSC_H vs width FSC_W (left column), FSC_H vs mCherry signal (middle column). Density plots show the distribution of the mCherry signal on a log scale with the blue line indicating the cutoff for mCherry positibe vs negative.

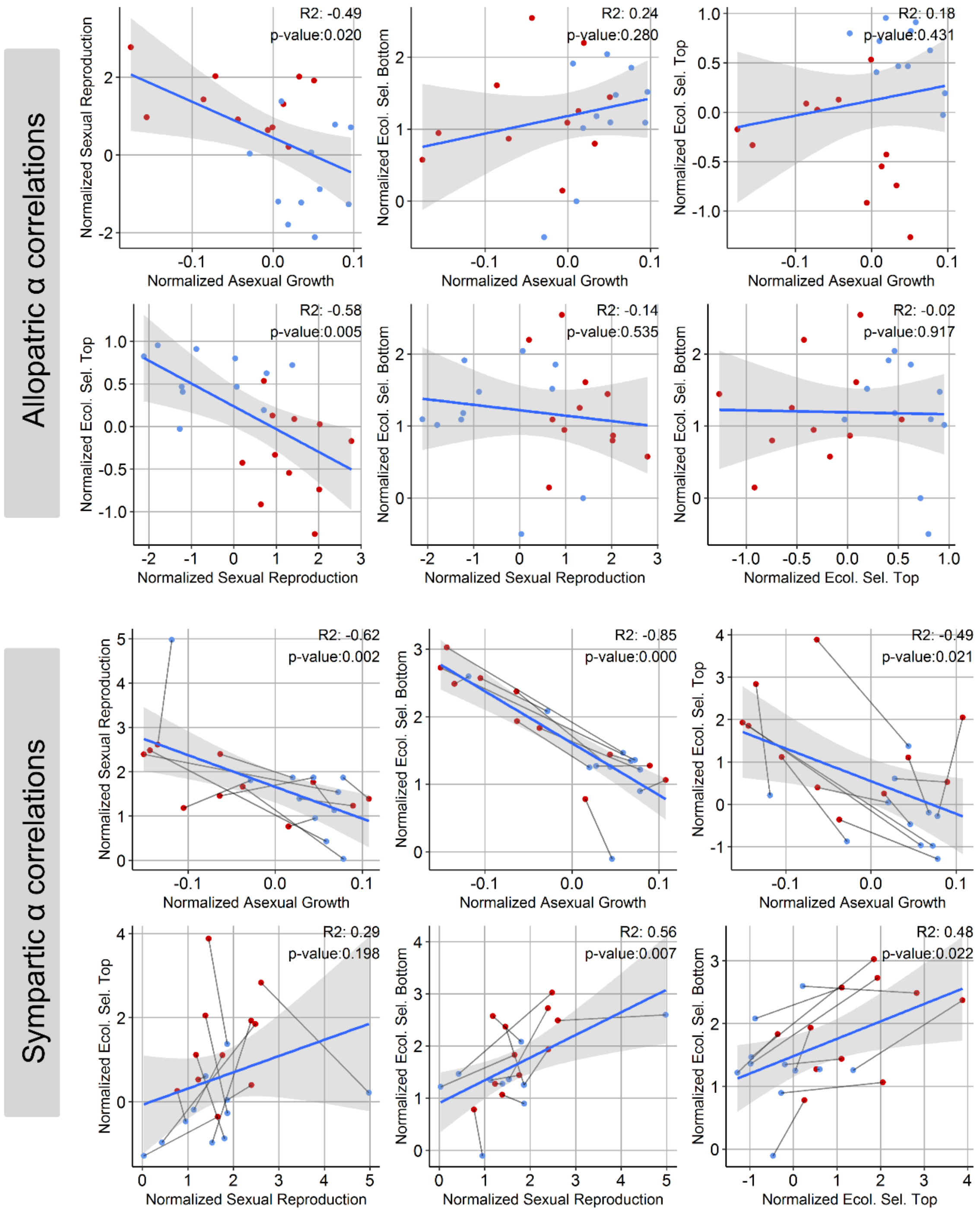


**Supplementary Figure 29.**

**Correlation between z-score standardized fitness components.** Example of pair-wise correlation estimates between fitness components, per treatment. In this example, allopatric and sympatric populations from the α genetic background are shown. In each plot, each point represents the median per population from 7 technical replicates. Populations are differentiates between top (blue) and bottom (red) selection regime for allopatry, and the same for the top and bottom ecotypes in sympatry. Adjusted $R^2$ and significance of the correlation (p-value) are shown for each plot.

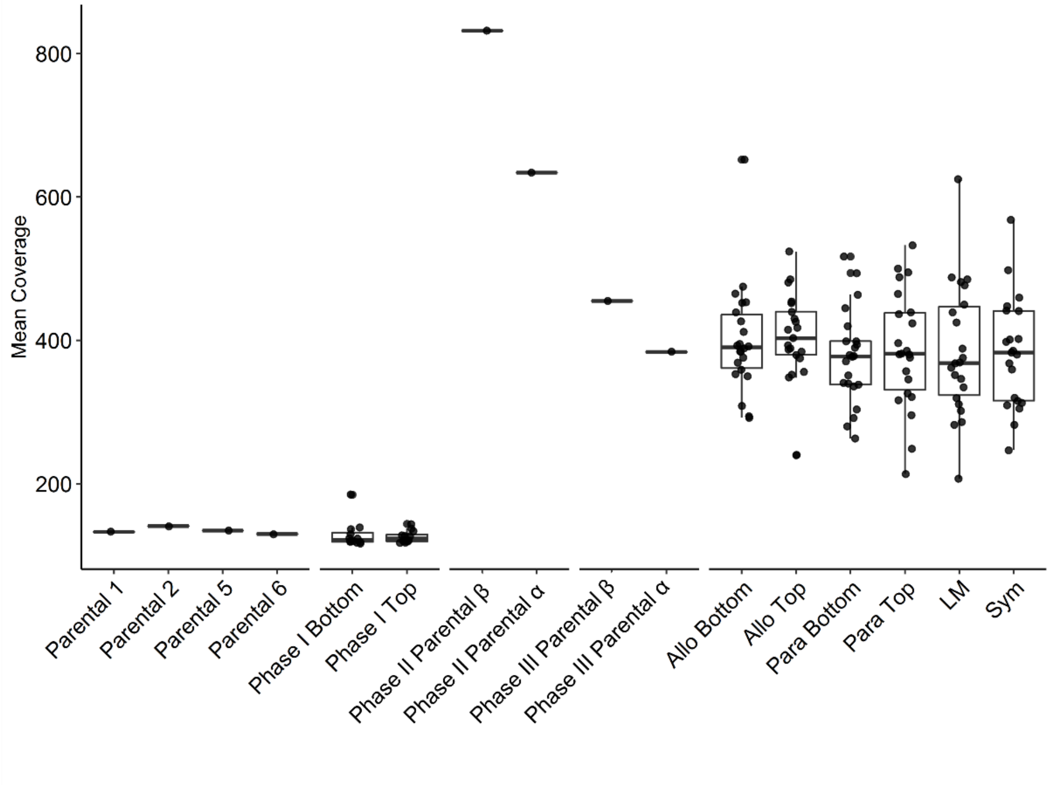


**Supplementary Figure 30.**

**Mean sequence coverage per population.** Box plot with mean coverage per sample. Coverage values are shown for all three phases of the experiment, including the four parental strains (**Supplementary Figure 5 -** Parentals P1, P2, P5, and P6), evolved strains from *phase* I (Phase I Top and Bottom), parentals of *phase II* and the ancestral populations for the two genetic backgrounds, and the different migration treatments of the experiment.

### Supplementary Tables

| | | α populations: | | | | | β populations: | | | | |
|---|---|---|---|---|---|---|---|---|---|---|---|
| | | Estimate | Std. Error | df | t value | P-Value | Estimate | Std. Error | df | t value | P-Value |
| Growth | Allopatry | 0.965 | 0.019 | 79.378 | 51.042 | **1.84E-62** | 0.946 | 0.021 | 63.545 | 44.127 | **2.18E-49** |
| | Local Mating | 0.081 | 0.027 | 79.378 | 3.019 | **3.41E-03** | 0.013 | 0.030 | 63.545 | 0.443 | 6.60E-01 |
| | Parapatry | 0.070 | 0.027 | 79.378 | 2.611 | **1.08E-02** | 0.065 | 0.030 | 63.545 | 2.131 | **3.69E-02** |
| | Sympatru | 0.000 | 0.027 | 79.378 | -0.014 | 9.89E-01 | -0.041 | 0.030 | 63.545 | -1.356 | 1.80E-01 |
| | Allo:FractionTop | 0.079 | 0.026 | 40.000 | 3.108 | **3.46E-03** | 0.020 | 0.021 | 40.000 | 0.951 | 3.47E-01 |
| | LM:FractionTop | -0.032 | 0.026 | 40.000 | -1.240 | 2.22E-01 | -0.043 | 0.021 | 40.000 | -2.015 | 5.07E-02 |
| | Para:FractionTop | 0.001 | 0.026 | 40.000 | 0.027 | 9.78E-01 | -0.073 | 0.021 | 40.000 | -3.445 | **1.36E-03** |
| | Sym:FractionTop | 0.069 | 0.026 | 40.000 | 2.715 | **9.72E-03** | 0.037 | 0.021 | 40.000 | 1.720 | 9.32E-02 |
| Mating | Allopatry | 1.356 | 0.298 | 79.998 | 4.551 | **1.88E-05** | 0.338 | 0.265 | 55.883 | 1.278 | 2.06E-01 |
| | Local Mating | -0.428 | 0.421 | 79.998 | -1.017 | 3.12E-01 | -0.054 | 0.374 | 55.883 | -0.145 | 8.85E-01 |
| | Parapatry | -0.574 | 0.421 | 79.998 | -1.361 | 1.77E-01 | -0.271 | 0.374 | 55.883 | -0.724 | 4.72E-01 |
| | Sympatru | 0.400 | 0.421 | 79.998 | 0.950 | 3.45E-01 | 0.316 | 0.374 | 55.883 | 0.844 | 4.02E-01 |
| | Allo:FractionTop | -1.858 | 0.420 | 40.172 | -4.419 | **7.33E-05** | -0.876 | 0.219 | 40.000 | -3.998 | **2.67E-04** |
| | LM:FractionTop | 0.215 | 0.420 | 40.172 | 0.511 | 6.12E-01 | 0.027 | 0.219 | 40.000 | 0.125 | 9.01E-01 |
| | Para:FractionTop | -0.552 | 0.420 | 40.172 | -1.313 | 1.97E-01 | -0.303 | 0.219 | 40.000 | -1.382 | 1.75E-01 |
| | Sym:FractionTop | -0.133 | 0.420 | 40.172 | -0.317 | 7.53E-01 | -0.326 | 0.219 | 40.000 | -1.487 | 1.45E-01 |
| TopSelection | Allopatry | -0.331 | 0.201 | 72.817 | -1.648 | 1.04E-01 | -0.187 | 0.183 | 76.852 | -1.022 | 3.10E-01 |
| | Local Mating | -0.051 | 0.284 | 72.817 | -0.181 | 8.57E-01 | 0.003 | 0.259 | 76.852 | 0.012 | 9.91E-01 |
| | Parapatry | 0.103 | 0.284 | 72.817 | 0.361 | 7.19E-01 | -0.182 | 0.259 | 76.852 | -0.704 | 4.83E-01 |
| | Sympatru | 1.747 | 0.284 | 72.817 | 6.150 | **3.78E-08** | 1.382 | 0.259 | 76.852 | 5.337 | **9.26E-07** |
| | Allo:FractionTop | 0.906 | 0.235 | 40.000 | 3.850 | **4.16E-04** | 0.975 | 0.231 | 40.000 | 4.216 | **1.38E-04** |
| | LM:FractionTop | 0.455 | 0.235 | 40.000 | 1.933 | 6.03E-02 | 0.462 | 0.231 | 40.000 | 2.000 | 5.23E-02 |
| | Para:FractionTop | -0.006 | 0.235 | 40.000 | -0.025 | 9.80E-01 | 0.476 | 0.231 | 40.000 | 2.060 | **4.60E-02** |
| | Sym:FractionTop | -1.673 | 0.235 | 40.000 | -7.109 | **1.32E-08** | -0.868 | 0.231 | 40.000 | -3.752 | **5.56E-04** |
| Bottom Selection | Allopatry | 1.223 | 0.194 | 70.680 | 6.310 | **2.14E-08** | -0.136 | 0.171 | 65.524 | -0.795 | 4.29E-01 |
| | Local Mating | 0.382 | 0.274 | 70.680 | 1.392 | 1.68E-01 | -0.156 | 0.242 | 65.524 | -0.644 | 5.22E-01 |
| | Parapatry | 0.033 | 0.274 | 70.680 | 0.121 | 9.04E-01 | -0.195 | 0.242 | 65.524 | -0.809 | 4.21E-01 |
| | Sympatru | 0.731 | 0.274 | 70.680 | 2.669 | **9.42E-03** | 0.139 | 0.242 | 65.524 | 0.576 | 5.67E-01 |
| | Allo:FractionTop | -0.071 | 0.219 | 40.000 | -0.325 | 7.47E-01 | -0.408 | 0.176 | 40.000 | -2.322 | **2.54E-02** |
| | LM:FractionTop | 0.232 | 0.219 | 40.000 | 1.062 | 2.95E-01 | -0.043 | 0.176 | 40.000 | -0.247 | 8.06E-01 |
| | Para:FractionTop | -0.042 | 0.219 | 40.000 | -0.191 | 8.50E-01 | -0.113 | 0.176 | 40.000 | -0.644 | 5.23E-01 |
| | Sym:FractionTop | -0.625 | 0.219 | 40.000 | -2.857 | **6.76E-03** | -0.428 | 0.176 | 40.000 | -2.436 | **1.94E-02** |

**Supplementary Table 1.**

**Results of a generalised linear model for each phenotype using median between technical replicates.** The model was calculated independently per fitness component and genetic background (α and β). In all cases, treatments were contrasted to the allopatric treatment. Selection regime (fraction top and bottom) was nested within treatment. Allo: allopatry, Para: parapatry, LM: Local Mating, Sym: Sympatry, Std. Error: Standard error, df: degrees of freedom, p values lower than 0.05 are shown in bold font.

| | | α populations: | | | | | β populations: | | | | |
|---|---|---|---|---|---|---|---|---|---|---|---|
| | | Estimate | Std. Error | df | t value | P-Value | Estimate | Std. Error | df | t value | P-Value |
| Growth | Allopatry | 0.969 | 0.018 | 568.000 | 55.199 | **1.98E-230** | 0.962 | 0.021 | 566.000 | 46.519 | **1.55E-195** |
| | Local Mating | 0.074 | 0.025 | 40.000 | 2.974 | **4.96E-03** | 0.009 | 0.029 | 40.000 | 0.320 | 7.51E-01 |
| | Parapatry | 0.077 | 0.025 | 40.000 | 3.110 | **3.44E-03** | 0.056 | 0.029 | 40.000 | 1.930 | 6.07E-02 |
| | Sympatry | -0.002 | 0.025 | 40.000 | -0.088 | 9.30E-01 | -0.041 | 0.029 | 40.000 | -1.386 | 1.73E-01 |
| | Allo:FractionTop | 0.078 | 0.019 | 568.000 | 4.195 | **3.17E-05** | 0.011 | 0.014 | 566.000 | 0.788 | 4.31E-01 |
| | LM:FractionTop | -0.038 | 0.019 | 568.000 | -2.026 | **4.32E-02** | -0.048 | 0.014 | 566.000 | -3.425 | **6.60E-04** |
| | Para:FractionTop | -0.007 | 0.019 | 568.000 | -0.390 | 6.97E-01 | -0.067 | 0.014 | 566.000 | -4.716 | **3.03E-06** |
| | Sym:FractionTop | 0.071 | 0.019 | 568.000 | 3.782 | **1.72E-04** | 0.028 | 0.014 | 566.000 | 1.995 | **4.66E-02** |
| Mating | Allopatry | 1.053 | 0.226 | 299.000 | 4.667 | **4.61E-06** | 0.251 | 0.248 | 302.000 | 1.014 | 3.11E-01 |
| | Local Mating | -0.240 | 0.321 | 40.000 | -0.745 | 4.61E-01 | -0.068 | 0.350 | 40.000 | -0.194 | 8.47E-01 |
| | Parapatry | -0.431 | 0.320 | 40.000 | -1.346 | 1.86E-01 | -0.279 | 0.350 | 40.000 | -0.797 | 4.30E-01 |
| | Sympatry | 0.622 | 0.319 | 40.000 | 1.950 | 5.82E-02 | 0.381 | 0.350 | 40.000 | 1.088 | 2.83E-01 |
| | Allo:FractionTop | -1.669 | 0.193 | 299.000 | -8.646 | **3.30E-16** | -0.868 | 0.119 | 302.000 | -7.294 | **2.66E-12** |
| | LM:FractionTop | 0.251 | 0.198 | 299.000 | 1.264 | 2.07E-01 | 0.037 | 0.120 | 302.000 | 0.307 | 7.59E-01 |
| | Para:FractionTop | -0.470 | 0.194 | 299.000 | -2.418 | **1.62E-02** | -0.312 | 0.119 | 302.000 | -2.624 | **9.12E-03** |
| | Sym:FractionTop | -0.169 | 0.193 | 299.000 | -0.877 | 3.81E-01 | -0.405 | 0.120 | 302.000 | -3.380 | **8.21E-04** |
| TopSelection | Allopatry | -0.281 | 0.176 | 535.000 | -1.593 | 1.12E-01 | -0.193 | 0.163 | 542.000 | -1.185 | 2.36E-01 |
| | Local Mating | -0.106 | 0.248 | 40.000 | -0.428 | 6.71E-01 | 0.090 | 0.230 | 40.000 | 0.392 | 6.97E-01 |
| | Parapatry | 0.023 | 0.249 | 40.000 | 0.093 | 9.26E-01 | -0.227 | 0.230 | 40.000 | -0.986 | 3.30E-01 |
| | Sympatry | 1.573 | 0.250 | 40.000 | 6.289 | **1.85E-07** | 1.425 | 0.232 | 40.000 | 6.149 | **2.92E-07** |
| | Allo:FractionTop | 0.870 | 0.138 | 535.000 | 6.306 | **6.02E-10** | 0.967 | 0.150 | 542.000 | 6.465 | **2.26E-10** |
| | LM:FractionTop | 0.416 | 0.135 | 535.000 | 3.094 | **2.08E-03** | 0.465 | 0.149 | 542.000 | 3.117 | **1.93E-03** |
| | Para:FractionTop | -0.017 | 0.136 | 535.000 | -0.126 | 9.00E-01 | 0.506 | 0.149 | 542.000 | 3.407 | **7.04E-04** |
| | Sym:FractionTop | -1.575 | 0.142 | 535.000 | -11.085 | **7.59E-26** | -0.950 | 0.157 | 542.000 | -6.031 | **3.02E-09** |
| Bottom Selection | Allopatry | 1.225 | 0.168 | 559.000 | 7.284 | **1.11E-12** | -0.077 | 0.156 | 561.000 | -0.493 | 6.22E-01 |
| | Local Mating | 0.434 | 0.237 | 40.000 | 1.829 | 7.49E-02 | -0.097 | 0.220 | 40.000 | -0.440 | 6.62E-01 |
| | Parapatry | -0.024 | 0.237 | 40.000 | -0.101 | 9.20E-01 | -0.089 | 0.220 | 40.000 | -0.403 | 6.89E-01 |
| | Sympatry | 0.648 | 0.237 | 40.000 | 2.729 | **9.39E-03** | 0.087 | 0.220 | 40.000 | 0.397 | 6.93E-01 |
| | Allo:FractionTop | -0.039 | 0.120 | 559.000 | -0.325 | 7.46E-01 | -0.417 | 0.112 | 561.000 | -3.737 | **2.05E-04** |
| | LM:FractionTop | 0.172 | 0.117 | 559.000 | 1.465 | 1.44E-01 | -0.034 | 0.111 | 561.000 | -0.307 | 7.59E-01 |
| | Para:FractionTop | -0.004 | 0.117 | 559.000 | -0.038 | 9.70E-01 | -0.130 | 0.110 | 561.000 | -1.177 | 2.40E-01 |
| | Sym:FractionTop | -0.562 | 0.117 | 559.000 | -4.817 | **1.88E-06** | -0.299 | 0.110 | 561.000 | -2.704 | **7.05E-03** |

**Supplementary Table 2.**

**Results of a generalised linear model for each phenotype using technical replicate measurements.** The model was calculated independently per fitness component and genetic background (α and β). In all cases, treatments were contrasted to the allopatric treatment. Selection regime (fraction top and bottom) was nested within treatment. Allo: allopatry, Para: parapatry, LM: Local Mating, Sym: Sympatry, Std. Error: Standard error, df: degrees of freedom, p values lower than 0.05 are shown in bold font.

| GO biological process complete | Genes in reference | # Genes mutated | Expected | Fold Enrichment | +/- | Raw P-value | FDR |
|---|---|---|---|---|---|---|---|
| 4,6-pyruvylated galactose residue biosynthetic process | 4 | 4 | 0.11 | 37.04 | + | 2.95E-05 | 1.03E-02 |
| ∟ cellular polysaccharide metabolic process | 48 | 9 | 1.30 | 6.95 | + | 1.42E-05 | 6.76E-03 |
| ∟ polysaccharide metabolic process | 52 | 10 | 1.40 | 7.12 | + | 3.81E-06 | 2.85E-03 |
| ∟ carbohydrate metabolic process | 126 | 15 | 3.40 | 4.41 | + | 3.34E-06 | 2.91E-03 |
| ∟ cellular carbohydrate metabolic process | 81 | 12 | 2.19 | 5.49 | + | 4.54E-06 | 2.38E-03 |
| cell-cell adhesion involved in flocculation | 9 | 5 | 0.24 | 20.58 | + | 1.93E-05 | 7.77E-03 |
| ∟ flocculation | 10 | 5 | 0.27 | 18.52 | + | 2.84E-05 | 1.06E-02 |
| ∟ aggregation of unicellular organisms | 12 | 6 | 0.32 | 18.52 | + | 4.24E-06 | 2.46E-03 |
| ∟ cell aggregation | 12 | 6 | 0.32 | 18.52 | + | 4.24E-06 | 2.77E-03 |
| adhesion between unicellular organisms via cell-wall interaction | 9 | 5 | 0.24 | 20.58 | + | 1.93E-05 | 8.42E-03 |
| ∟ adhesion between unicellular organisms | 11 | 6 | 0.30 | 20.21 | + | 2.89E-06 | 3.02E-03 |
| ∟ multi organism cell adhesion | 11 | 6 | 0.30 | 20.21 | + | 2.89E-06 | 3.78E-03 |
| ∟ cell adhesion | 15 | 8 | 0.40 | 19.76 | + | 6.54E-08 | 3.42E-04 |
| ∟ biological adhesion | 15 | 8 | 0.40 | 19.76 | + | 6.54E-08 | 1.71E-04 |
| ∟ cell-cell adhesion | 12 | 7 | 0.32 | 21.61 | + | 2.88E-07 | 5.03E-04 |
| polysaccharide catabolic process | 13 | 5 | 0.35 | 14.25 | + | 7.59E-05 | 2.34E-02 |
| fungal-type cell wall biogenesis | 54 | 8 | 1.46 | 5.49 | + | 1.90E-04 | 4.97E-02 |
| ∟ cell wall organization or biogenesis | 97 | 11 | 2.62 | 4.20 | + | 1.08E-04 | 3.15E-02 |
| ∟ fungal-type cell wall organization or biogenesis | 86 | 11 | 2.32 | 4.74 | + | 4.00E-05 | 1.31E-02 |
| regulation of anatomical structure morphogenesis | 81 | 10 | 2.19 | 4.57 | + | 1.19E-04 | 3.27E-02 |

**Supplementary Table 3.**

GO-term analysis for the 138 mapped genes from the 140 genes that had multiple independent mutations using pombase GO Ontology database (Released 2019-12-09) with Fisher's exact test with False discovery rate to correct for multiple testing.
